# Phospholipid flippases and Sfk1 are essential for the retention of ergosterol in the plasma membrane

**DOI:** 10.1101/2020.10.19.346320

**Authors:** Takuma Kishimoto, Tetsuo Mioka, Eriko Itoh, David E. Williams, Raymond J. Andersen, Kazuma Tanaka

**Author notes:** Corresponding Author (TK), (KT).

## Abstract

Sterols are important lipid components of the plasma membrane (PM) in eukaryotic cells, but it is unknown how the PM retains sterols at a high concentration. Phospholipids are asymmetrically distributed in the PM, and phospholipid flippases play an important role in generating this phospholipid asymmetry. Here, we provide evidence that phospholipid flippases are essential for retaining ergosterol in the PM of yeast. A mutant in three flippases, Dnf1-Lem3, Dnf2-Lem3, and Dnf3-Crf1, and a membrane protein, Sfk1, showed a severe growth defect. We recently identified Sfk1 as a PM protein involved in phospholipid asymmetry. The PM of this mutant showed high permeability and low density, and many nutrient transporters failed to localize to the PM. Staining with the sterol probe filipin and the expression of a sterol biosensor revealed that ergosterol was not retained in the PM. Instead, ergosterol accumulated in an esterified form in lipid droplets. We propose that ergosterol is retained in the PM by the asymmetrical distribution of phospholipids and the action of Sfk1. Once phospholipid asymmetry is severely disrupted, sterols might be exposed on the cytoplasmic leaflet of the PM and actively transported to the endoplasmic reticulum by sterol transfer proteins.

## Introduction

Heterogeneity in the distribution of membrane phospholipids and sterols is essential for the diverse functions of cells. In the plasma membrane (PM) of eukaryotic cells, phosphatidylcholine (PC), sphingolipids, and gangliosides are predominantly distributed in the extracellular leaflet, whereas phosphatidylethanolamine (PE), phosphatidylserine (PS), and other charged lipids are mainly localized to the cytoplasmic leaflet [1–3]. This asymmetric distribution of phospholipids is controlled by three types of lipid translocators: flippase, catalyzing inward phospholipid translocation (flip) [4–6]; floppase, catalyzing outwards phospholipid translocation (flop) [5, 7, 8]; and scramblase, catalyzing bidirectional phospholipid translocation [9].

Accumulating genetic and biochemical evidence indicates that flippases are integrally linked to phospholipid asymmetry of the organelle membrane from yeast to mammalian cells. Flippases, which are type 4 P-type ATPases (P4-ATPases), have the ability to translocate phospholipids from the extracellular leaflet of the PM or luminal leaflet of endomembranes to the cytoplasmic leaflet [6]. At the cellular level, flippases are associated with diverse physiological functions. Flippases in endomembranes function primarily in membrane trafficking processes [10–18], whereas those located in the PM are involved in multiple cellular processes: membrane trafficking [10, 12, 13, 19], apoptosis signaling [20], mating signaling [21], the apical membrane barrier [22], cell polarity [23–25], and cell migration [26].

Flippases form heterodimeric complexes with noncatalytic subunits of the Cdc50 family. Budding yeast has five P4-ATPases: Drs2, Dnf1, Dnf2, Dnf3, and Neo1 [27] and three Cdc50 family member proteins: Cdc50, Lem3, and Crf1 [13, 14]. Drs2 and Dnf3 interact with Cdc50 and Crf1, respectively, and are mainly localized to the endomembrane, such as the *trans*-Golgi network (TGN) and endosomes. On the other hand, both Dnf1 and Dnf2 form complexes with Lem3 and are mainly localized to the PM [12, 28]. Except for Neo1, interactions between the P4-ATPases and Cdc50 subunits are essential for endoplasmic reticulum (ER) exit and proper subcellular localization of the complexes but may also contribute to their lipid translocase activity and functions [13, 14, 29–32]. Thus, phenotypes in P4-ATPase mutants are phenocopied by their subunit mutants [13, 14].

Dnf1/2-Lem3 complexes are endocytosed but recycled back to the PM through the endocytic recycling pathway [13, 14], maintaining the localization of these complexes to the PM. Genetic analyses suggested that the Dnf1/2-Lem3 complexes have PE and PS translocation activity [12, 28, 33, 34]. Considering the localization and activity of Dnf1/2-Lem3 complexes, they maintain phospholipid asymmetry predominantly at the PM. Compared to the other four P4-ATPases, little is known about the activity and function of the Dnf3-Crf1 complex. However, the deletion of *DNF3* increases the sensitivity of the *dnf1*Δ *dnf2*Δ double mutant to the PE-binding peptide duramycin [21], and Dnf3 is implicated in the translocation of PS across the PM [35], suggesting possible functions of the Dnf3-Crf1 complex in PM phospholipid translocation.

In addition to Dnf1/2-Lem3, some regulators are involved in phospholipid asymmetry of the PM. Serine/threonine kinases Fpk1/2 upregulate Dnf1/2 flippase activity via phosphorylation [36]. Pdr5p and Yor1p, two multidrug ABC transporters [12, 37], and Opt2, a member of the oligopeptide transporter family [38], are implicated in the flop of phospholipids. Recently, we isolated Sfk1 as a multicopy suppressor of the *lem3*Δ mutant; overexpression of Sfk1 suppressed PE and PS exposure in the PM [39]. Sfk1 is a conserved transmembrane protein belonging to the TMEM150/FRAG1/DRAM family [40]. From genetic analyses, we proposed that Sfk1 might negatively regulate the transbilayer movement of phospholipids irrespective of direction in an unprecedented way. The *lem3*Δ *sfk1*Δ double mutant exhibits more severe defects in PE and PS asymmetry in the PM than the *lem3*Δ mutant, and the *lem3*Δ *sfk1*Δ mutant exhibits increased permeability of the PM [39]. However, these mutations do not affect cell growth. Given that PM phospholipid asymmetry is commonly observed in eukaryotes, it may be speculated that phospholipid asymmetry plays an important role (e.g., is essential for cell growth). Thus, there might be a gene that functions redundantly with *LEM3* and *SFK1* to control phospholipid asymmetry.

Another important feature of the PM is that this membrane is rich in sterols. Sterols such as mammalian cholesterol and the fungal ergosterol are essential membrane components with tightly controlled homeostasis [41]. At the cellular level, the PM contains approximately 30-40 mol% cholesterol in PM lipids, whereas ER contains approximately 5 mol% cholesterol [42, 43]. Sterols are inserted into lipid membranes through the interaction between 3-hydroxyl groups and hydrocarbon rings of sterols and polar head groups and hydrocarbon chains of phospholipids, respectively [44]. Each phospholipid has a different affinity for sterols, which determines the strength of their interaction with sterols [45, 46]. Sphingolipids, PC, and PS interact strongly with sterol, whereas phospholipids with small polar head groups and unsaturated fatty acyl tails exhibit weaker interactions [47–49]. Numerous studies have suggested that these interactions contribute to the properties of the PM, including tight packing, high rigidity, and low permeability. However, it is unclear how the PM retains such a high concentration of sterols and whether the asymmetric distribution of PE and PS is involved in retaining sterols in the PM.

In this study, we searched for genes that functionally interact with *LEM3* and *SFK1* by synthetic lethal genetic screening and identified *dnf3* and *crf1* as interacting partners. The conditional *crf1 lem3 sfk1* triple mutant cannot maintain ergosterol in the PM and instead accumulates esterified ergosterol in the lipid droplet (LD). Our results suggest that Dnf1/2-Lem3 and Dnf3-Crf1 flippases and Sfk1 function cooperatively to maintain the phospholipid asymmetry of the PM, which is essential for sterol retention in the PM and thus for the homeostatic control of sterol.

## Results

### Dnf3-Crf1 flippase is involved in PM phospholipid asymmetry together with Dnf1/2-Lem3 flippases and Sfk1

To isolate genes involved in the regulation of phospholipid asymmetry of the PM in conjunction with Lem3 and Sfk1, we searched for mutations that display synthetic lethality with *lem3*Δ *sfk1*Δ mutations at 30°C. We isolated a new allele of the flippase noncatalytic subunit *crf1*Δ (Fig 1A). To confirm this synthetic lethality, we crossed the *crf1*Δ *lem3*Δ mutant to the *lem3*Δ *sfk1*Δ mutant, followed by tetrad analysis (Fig 1B). The *crf1Δ lem3*Δ *sfk1*Δ triple mutant did not germinate at 30°C but germinated at 25°C despite severe growth defects (Fig 1B), which allowed us to obtain the *crf1*Δ *lem3*Δ *sfk1*Δ triple mutant for further analysis. We next tested the growth of the *crf1*Δ *lem3*Δ *sfk1*Δ triple mutant at 30 and 37°C. The triple mutant grew very slowly at 30°C and showed lethality at 37°C (Fig 1C). The deletion of *DNF3*, which encodes the catalytic subunit of Crf1 [13], also grew poorly when combined with *lem3*Δ *sfk1*Δ (Fig 1D).

**Fig 1.**
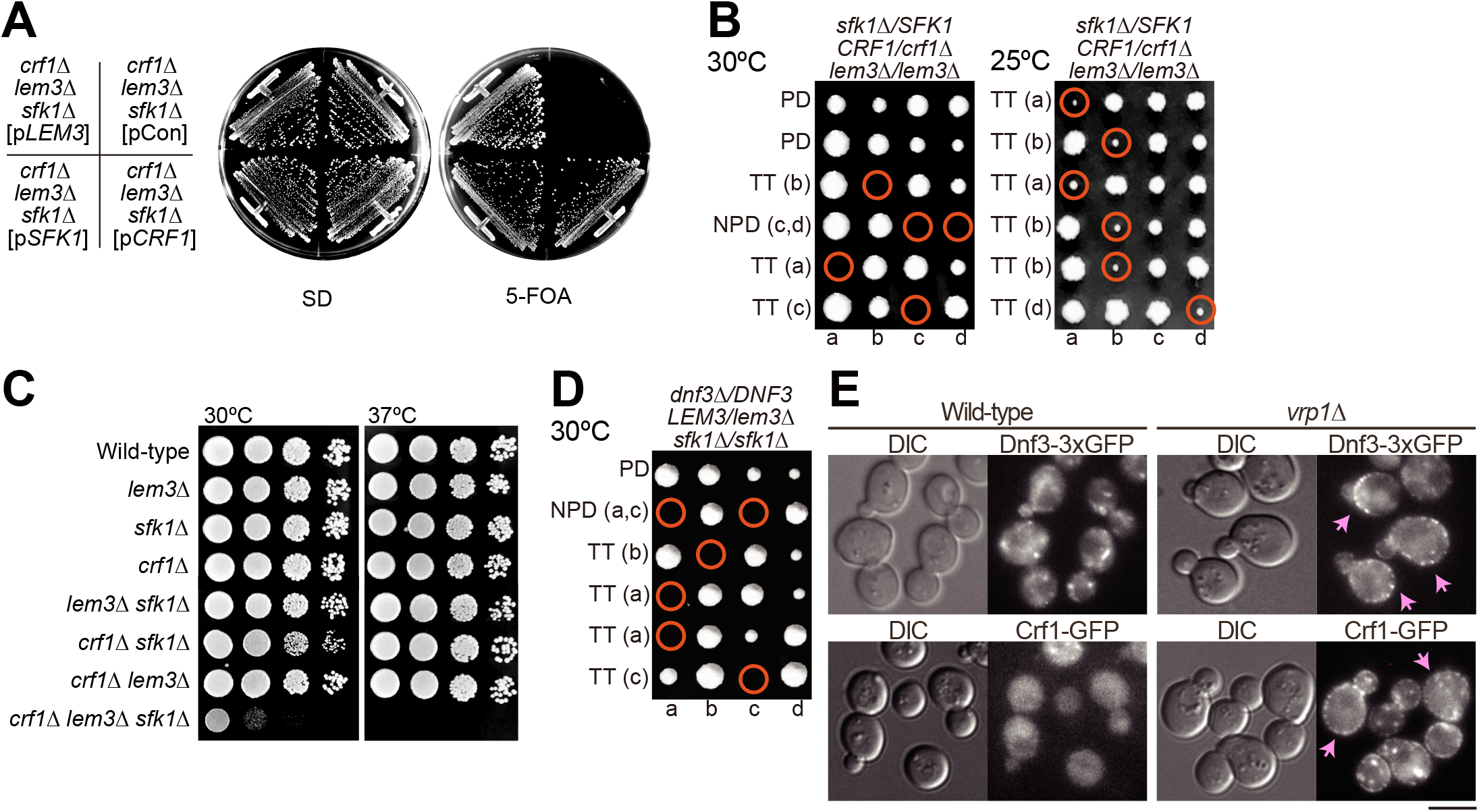
Synthetic growth defects of the *crf1 lem3 sfk1* mutant. (A) Growth profiles on 5-fluoroorotic acid (5-FOA) plate medium. The *crf1Δ lem3*Δ *sfk1*Δ mutant harboring pRS316-*SFK1* was transformed with YCplac111 (pCon), YCplac111-*LEM3* (p*LEM3*), YCplac111-*CRF1* (p*CRF1*), or pRS315-*SFK1* (p*SFK1*). Transformants were streaked onto an SD-Leu + 5-FOA plate and grown at 30°C for 3 d. The cells that require pRS316-*SFK1* for growth are sensitive to 5-FOA because pRS316 contains the *URA3* gene [51]. (B) Tetrad dissection analysis. Diploid cells with the indicated genotype were sporulated, dissected, and grown at either 30 or 25°C for 4 d. Colonies were replica-plated onto selective media to determine the segregation of the marked mutant alleles. Tetrad genotypes (TT, tetratype; PD, parental ditype; and NPD, nonparental ditype) are indicated, and the identities of the triple mutant segregants are shown in parentheses (red circles). (C) Growth profiles by spot growth assay. As described in the “Materials and Methods”, tenfold serial dilutions of cell cultures were spotted onto YPDA and grown for 1.5 d at 30 or 37°C. (D) Synthetic growth defects of the *dnf3*Δ *lem3*Δ *sfk1*Δ mutant. Tetrad analysis was performed as in (B). (E) Localizations of Dnf3-3xGFP (triple tandem GFP) and Crf1-GFP in the endocytosis defective *vrp1*Δ mutant. Cells were grown to mid-log phase in YPDA medium at 30°C. Arrows indicate the cells showing the PM localization of examined proteins. Bar, 5 µm. DIC, differential interference contrast.

Dnf3 is mainly localized to endosomal/Golgi membranes [10, 12], but it was suggested that Dnf3 also functions at the PM [35]. To examine whether Dnf3-Crf1 is transported to the PM, we used the endocytosis-deficient *vrp1*Δ mutant [50]. Both Dnf3-3xGFP (triple tandem green fluorescent protein [GFP]) and Crf1-GFP were localized to intracellular structures but were barely detectable in the PM of the wild-type. However, they were observed in the PM when endocytosis was inhibited (Fig 1E). This result indicates that the Dnf3-Crf1 flippase is transported between the PM and endomembranes, similar to Drs2-Cdc50 [14]. These results raise the possibility that the synthetic growth defect of the *crf1*Δ *lem3*Δ *sfk1*Δ mutant is caused by defects in the PM, and this point was analyzed further.

We attempted to perform phenotypic analysis of the *crf1*Δ *lem3*Δ *sfk1*Δ triple mutant. However, the expression of some GFP-fused proteins resulted in lethality in the *crf1*Δ *lem3*Δ *sfk1*Δ background. Thus, we constructed temperature-sensitive (ts) mutants of *SFK1* by random mutagenesis in the *lem3*Δ *crf1*Δ background as described in the “Materials and Methods”. The *crf1 lem3 sfk1-2* mutant exhibited acceptable growth at 30°C but a severe growth defect at 37°C (Fig 2A). From the growth profiles of the *crf1*Δ *lem3*Δ *sfk1-2* mutant at 30 and 37°C (S1 Fig), we determined that phenotypes of the triple mutant were analyzed after culturing for 6 h after the shift to 37°C. DNA sequencing of the *sfk1-2* mutant allele revealed that *sfk1-2* contained one mutation that resulted in an amino acid substitution W16R (Fig 2B), which was located in the N-terminal transmembrane region. Wild-type Sfk1-3xGFP was localized to the PM, whereas the mutant Sfk1-2-3xGFP exhibited a lower signal in the PM and the ER at 30°C and was barely detectable at 37°C (Fig 2C).

**Fig 2.**
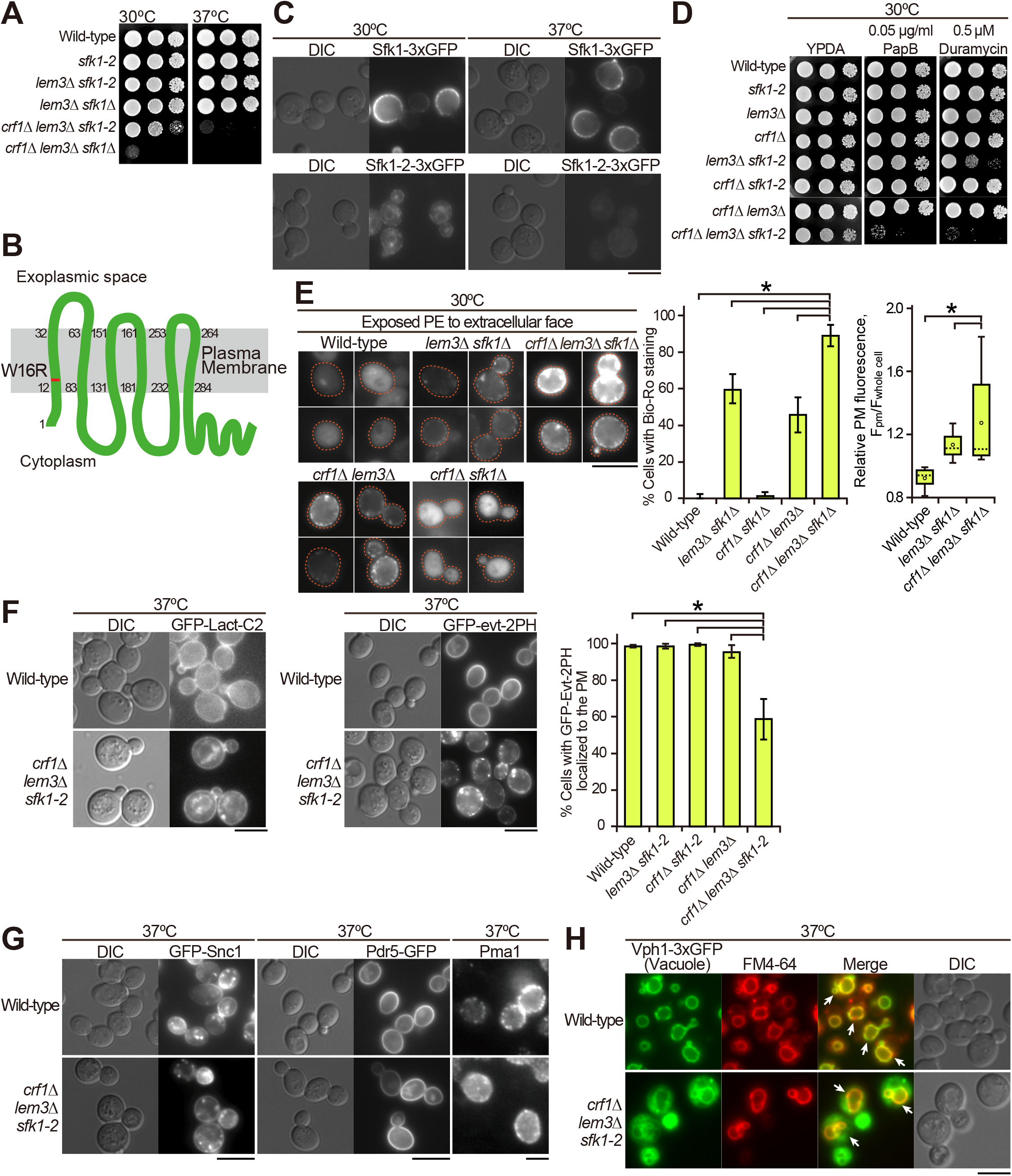
The *crf1*Δ *lem3*Δ *sfk1* triple mutants show severe defects in phospholipid asymmetry but not in membrane trafficking. (A) Isolation of the *sfk1-2* ts mutant. Tenfold serial dilutions of cell cultures were spotted onto a YPDA plate, followed by incubation at 30 or 37°C for 1.5 d. (B) Amino acid substitution of the Sfk1-2 mutant protein. The W16R substitution occurs in the first transmembrane domain of Sfk1-2. (C) Localization of the Sfk1-2 mutant protein fused with 3xGFP. Cells were grown in YPDA medium to mid-log phase at 30°C and then shifted to 37°C, followed by incubation for 6 h. (D) The *crf1*Δ *lem3*Δ *sfk1-2* triple mutant was sensitive to PapB and duramycin. Tenfold serial dilutions were spotted onto a YPDA plate containing PapB or duramycin, followed by incubation at 30°C for 2 d. (E) PE was most exposed in the *crf1*Δ *lem3*Δ *sfk1*Δ mutant. *Left panels*: cells were cultured in YPDA at 30°C, and exposed PE was visualized by staining with Bio-Ro and Alexa Fluor 488–labeled streptavidin. Dashed lines indicate cell edges. *Middle panel*: the percentages of cells showing PE exposure were determined and are expressed as the mean ± standard deviation (S.D.) of three independent experiments (n > 81 cells in total for each strain). An asterisk indicates a significant difference, as determined by the Tukey–Kramer test (p < 0.05). *Right panel*: fluorescence intensity at the PM was quantitated as described in the “Materials and Methods”. The ratio of the fluorescence at the PM (F_pm_)/that of whole cell (F_whole cell_) was determined and expressed in a boxplot (whiskers: maximum and minimum values; box: first quartile, median, and third quartile; circle: average). The numbers of cells analyzed were 23, 38, and 35 for the wild-type, *lem3 sfk1*Δ, and *crf1*Δ *lem3*Δ *sfk1*, respectively. An asterisk indicates a significant difference, as determined by the Tukey–Kramer test (p < 0.05). (F) GFP-evt-2PH was mislocalized in the *crf1*Δ *lem3*Δ *sfk1-2* mutant. Cells were cultured as in Fig 2C. *Right panel*: the percentage of cells with GFP-evt-2PH at the PM was determined and is expressed as the mean ± S.D. of three independent experiments (n >154 cells in total for each strain). An asterisk indicates a significant difference, as determined by the Tukey–Kramer test (p < 0.05). (G) Normal localization of PM proteins in the *crf1*Δ *lem3*Δ *sfk1-2* mutant. Cells were cultured as in Fig 2C. Pma1 was detected by immunostaining as described in the “Materials and Methods”. (H) Endocytosis was not significantly affected in the *crf1*Δ *lem3*Δ *sfk1-2* mutant. Cells expressing the vacuole membrane marker Vph1-3xGFP were cultured as in Fig 2C. Then, cells were incubated with FM4-64 on ice for 30 min, followed by incubation at 37°C for 30 min. Arrows indicate the colocalization of FM4-64 and Vph1-3xGFP. Bars, 5 µm.

Phospholipid asymmetry defects cause the exposure of PS and PE to the extracellular leaflet of the PM. The exposed PS and PE can be indirectly measured by examination of the growth sensitivities of the mutants to the PS-binding cyclodepsipeptide papuamide B (PapB) and PE-binding tetracyclic peptide duramycin. We previously reported that the *lem3*Δ *sfk1*Δ double mutant exhibited high sensitivities to both peptides [39]. Thus, we first tested the growth sensitivity of the *crf1*Δ *lem3*Δ *sfk1-2* triple mutant to these peptides at 30°C (Fig 2D). The addition of either the *crf1*Δ or *dnf3*Δ mutation to the *lem3* mutant elevated the sensitivities to both peptides (S2A Fig), consistent with a previous report on the *dnf1*Δ *dnf2*Δ *dnf3*Δ mutant [21]. The *crf1*Δ *lem3*Δ *sfk1-2* mutant did not grow at the concentrations at which the *crf1*Δ *lem3*Δ and *lem3*Δ *sfk1-2* double mutants could grow (Fig 2D), suggesting that the *crf1*Δ *lem3*Δ *sfk1-2* triple mutant exposed more PS and PE even at the permissive temperature than did the double mutants. To further confirm the defect in phospholipid asymmetry in the triple mutant, we next visualized the PE exposed to the extracellular surface using the PE-binding biotinylated Ro 09-0198 peptide (Bio-Ro). Fluorescence signals were not detected in either the wild-type or *crf1*Δ *sfk1*Δ double mutant but were detected in both the *crf1*Δ *lem3* (45%) and *lem3*Δ *sfk1*Δ double mutants (58%) (Fig 2E, left and middle panels). In the *crf1*Δ *lem3*Δ *sfk1*Δ triple mutant, the proportion of cells with fluorescent signals increased to 85%. Furthermore, the average signal intensity in the triple mutant was 1.35-fold higher than that in the *lem3*Δ *sfk1*Δ mutant (Fig 2E, right panel).

Next, we examined PS distribution in the cytoplasmic leaflet of the PM in the *crf1*Δ *lem3*Δ *sfk1-2* triple mutant. To visualize PS, we expressed PS biosensors, the C2 domain of lactadherin (Lact-C2) [52] and the pleckstrin homology (PH) domain of evectin-2 (evt-2PH) [53]. GFP-Lact-C2 was mainly distributed in the PM of the examined cells, but intracellular localization was also observed in the triple mutant (Fig 2F, left panel, S2B Fig). In contrast, GFP-evt-2PH was normally distributed only to the PM in the wild-type and double mutants (more than 96% of cells with PM distribution), whereas the GFP-evt-2PH signal was lost or significantly reduced from the PM in the *crf1*Δ *lem3*Δ *sfk1-2* triple mutant (59% of cells with PM distribution) (Fig 2F, middle and right panels, S2C Fig). We speculate that Lact-C2 has a higher affinity for PS, resulting in the detection of a lower level of PS at the PM. Taken together, these results suggest that the asymmetric distribution of PE and PS was most disturbed in the *crf1*Δ *lem3Δ sfk1* triple mutants.

As the Dnf3-Crf1 complex was mainly localized to endosomal/TGN compartments (Fig 1E) [10, 12], the *crf1*Δ *lem3*Δ *sfk1-2* mutant may exhibit a defect in membrane trafficking. We examined the localization of the endocytic recycling marker GFP-Snc1, which is mainly localized to polarized PM sites [54], but its localization was not affected in the *crf1*Δ *lem3*Δ *sfk1-2* triple mutant at 37°C (Fig 2G). Similarly, two PM proteins, Pdr5-GFP (ABC transporter) [55] and Pma1 (H^+^-ATPase) [56], were normally transported to the PM in the *crf1*Δ *lem3Δ sfk1-2* triple mutant (Fig 2G). We also examined endocytosis in the *crf1*Δ *lem3*Δ *sfk1-2* triple mutant by uptake of the lipophilic dye FM4-64 [57]. The FM4-64 signal was well colocalized to the vacuole membrane marker Vph1-3xGFP [58] in both the wild-type and *crf1*Δ *lem3*Δ *sfk1-2* triple mutant after 30 min of incubation, suggesting that the triple mutant did not have obvious defects in endocytosis (Fig 2H). These results suggest that the *crf1*Δ *lem3*Δ *sfk1-2* triple mutant is not defective in membrane trafficking to or from the PM.

### Multiple functions of the PM are impaired in the *crf1*Δ *lem3*Δ *sfk1* triple mutants

Phospholipid asymmetry defects may have a profound effect on PM functions. Previously, we showed that the *lem3*Δ *sfk1*Δ double mutant exhibits an increase in membrane permeability by measuring rhodamine dye uptake [39]. This experiment was performed in the *crf1*Δ *lem3*Δ *sfk1*Δ triple mutant, and the results suggest that the permeability is further enhanced in the triple mutant compared with that in the *lem3*Δ *sfk1*Δ double mutant (Fig 3A). The large increase in membrane permeability prompted us to examine whether the lipid composition changes in the PM of the *crf1*Δ *lem3*Δ *sfk1-2* triple mutant. We performed sucrose density gradient fractionation to isolate the PM. In wild-type, PM markers, both Pdr5-GFP [55] and Pma1 [59, 60], were recovered in high-density fractions, whereas Kex2, which is localized to endosomal/TGN compartments [54, 61], peaked at a lower density (Fig 3B). However, in the *crf1*Δ *lem3*Δ *sfk1-2* triple mutant, Pdr5-GFP, Pma1, and Kex2 were recovered together in lower density fractions (fractions 4-8) (Fig 3B). Pdr5-GFP and Pma1 were normally localized to the PM in the *crf1*Δ *lem3*Δ *sfk1-2* triple mutant in microscopic analysis (Fig 2G), suggesting a decrease in PM density, which makes PM isolation from the triple mutant technically challenging. Thus, we measured the phospholipid composition in the total cellular lipids. No significant difference in lipid composition was found in the double and triple mutants (S3 Fig).

**Fig 3.**
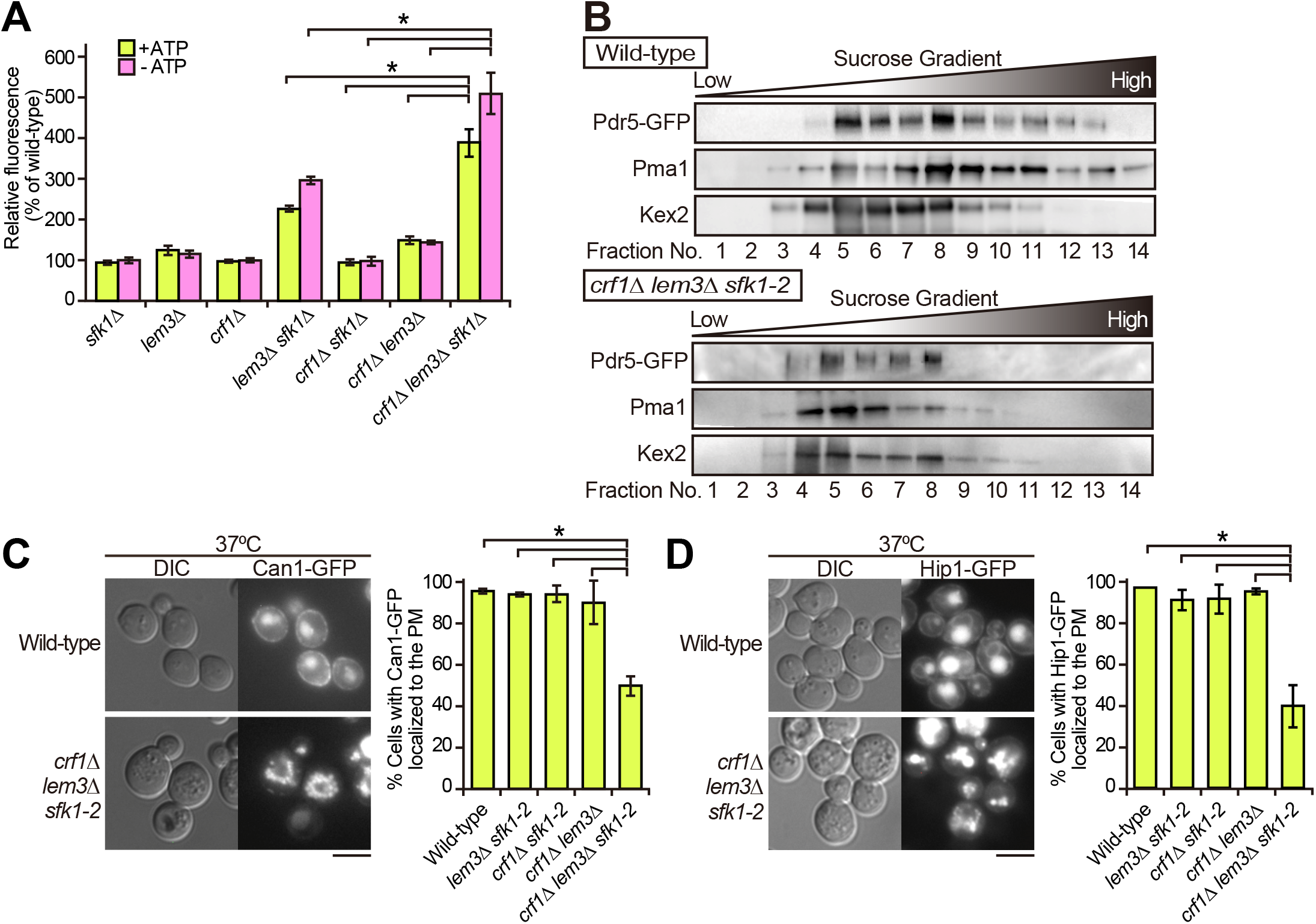
Multiple defects in the PM of the *crf1*Δ *lem3*Δ *sfk1* triple mutants. (A) Rhodamine uptake is increased in the *crf1*Δ *lem3*Δ *sfk1*Δ triple mutant. Cells were cultured in YPDA medium at 30°C, preincubated in SD medium with 1 mM sodium azide for 30 min at 30°C, and incubated with rhodamine 6G for 60 min at 30°C. Rhodamine accumulation was measured as described in the “Materials and Methods”. Values represent the mean ± S.D. from three independent experiments. Asterisks indicate a significant difference, as determined by the Tukey–Kramer test (p < 0.05). (B) Sucrose density gradient centrifugation analysis of PM proteins, Pdr5-GFP and Pma1, and a TGN/endosome protein, Kex2, in the *crf1*Δ *lem3*Δ *sfk1-2* triple mutant. Cells were cultured as in Fig 2C. Cell lysates were prepared from the wild-type and the *crf1*Δ *lem3*Δ *sfk1-2* triple mutant expressing Pdr5-GFP and fractionated in 27–60% sucrose step density gradients as described in the “Materials and Methods”. Equivalent volumes from each fraction were subjected to sodium dodecyl sulfate polyacrylamide gel electrophoresis (SDS-PAGE), and proteins were detected by immunoblotting. (C and D) The distributions of GFP-fused amino acid transporters, Can1-GFP (C) and Hip1-GFP (D), were analyzed in the wild-type and the *crf1*Δ *lem3*Δ *sfk1-2* triple mutant. Cells were cultured as in Fig 2C. Bar, 5 µm. *Right panels*: the percentage of cells with Can1-GFP or Hip1-GFP at the PM was determined and is expressed as the mean ± S.D. of three independent experiments (n>170 cells in total for each strain). An asterisk indicates a significant difference, as determined by the Tukey–Kramer test (p < 0.05).

The PM integrity defects may impact critical functions of the PM, such as localization of PM transporters involved in nutrient uptake. We analyzed the localization of PM transporters in the *crf1*Δ *lem3*Δ *sfk1-2* mutant. In the wild-type and double mutants, incubation at 37°C had little effect on the PM localization of amino acid transporters Can1-GFP and Hip1-GFP [56, 62]; more than 94% of the cells displayed PM localization (Fig 3C and 3D, S4A Fig). However, both transporters were not localized to the PM and instead were only localized to the vacuole in many of the *crf1*Δ *lem3*Δ *sfk1-2* mutant cells; the percentage of Can1-GFP and Hip1-GFP at the PM decreased to 58% and 41%, respectively (Fig 3C and 3D). Other amino acid transporters (Alp1, Lyp1, Tat1, and Ptr2) and hexose transporters (Hxt2-4) [62–65] also failed to localize to the PM in the *crf1*Δ *lem3*Δ *sfk1-2* mutant (S4B Fig). To test whether these transporters once reached the PM before being transported to the vacuole, we next examined Can1-GFP and Hip1-GFP localizations in cells treated with Latrunculin-A (LAT-A), which inhibits actin-dependent endocytic internalization by interfering with actin patch assembly [66]. In the triple mutant, LAT-A treatment increased the number of cells showing PM localization of Can1-GFP (67%) and Hip1-GFP (63%) compared with the control DMSO treatment (Can1-GFP, 32%; Hip1-GFP, 38%) (S4C Fig). We conclude that the *crf1*Δ *lem3*Δ *sfk1* triple mutants cannot maintain PM integrity, causing multiple defects in PM function.

### Isolation of *KES1* as a multicopy suppressor of the *crf1*Δ *lem3*Δ *sfk1-2* mutation

To explore the essential functions of phospholipid asymmetry in the PM, we screened for multicopy suppressors of the ts growth defect of the *crf1*Δ *lem3*Δ *sfk1-2* triple mutant. We found that overexpression of *KES1* suppressed the growth defect (Fig 4A). Kes1, also known as Osh4, is an oxysterol-binding protein (OSBP) homolog (Osh) that is implicated in sterol transport within cells [67]. Budding yeast contains seven Osh homologs, Osh1-7, that exchange specific lipids between organelles [68]. We next tested whether overexpression of other Osh proteins, except for *OSH1*, which localizes to the nucleus-vacuole junction [69], could suppress the growth defect of the *crf1*Δ *lem3*Δ *sfk1-2* triple mutant. Only *KES1* overexpression suppressed the ts growth defect of the *crf1*Δ *lem3*Δ *sfk1-2* triple mutant (Fig 4A). Increased rhodamine uptake and Can1-GFP mislocalization were also suppressed by *KES1* overexpression in the *crf1*Δ *lem3*Δ *sfk1-2* triple mutant (Fig 4B and 4C).

**Fig 4.**
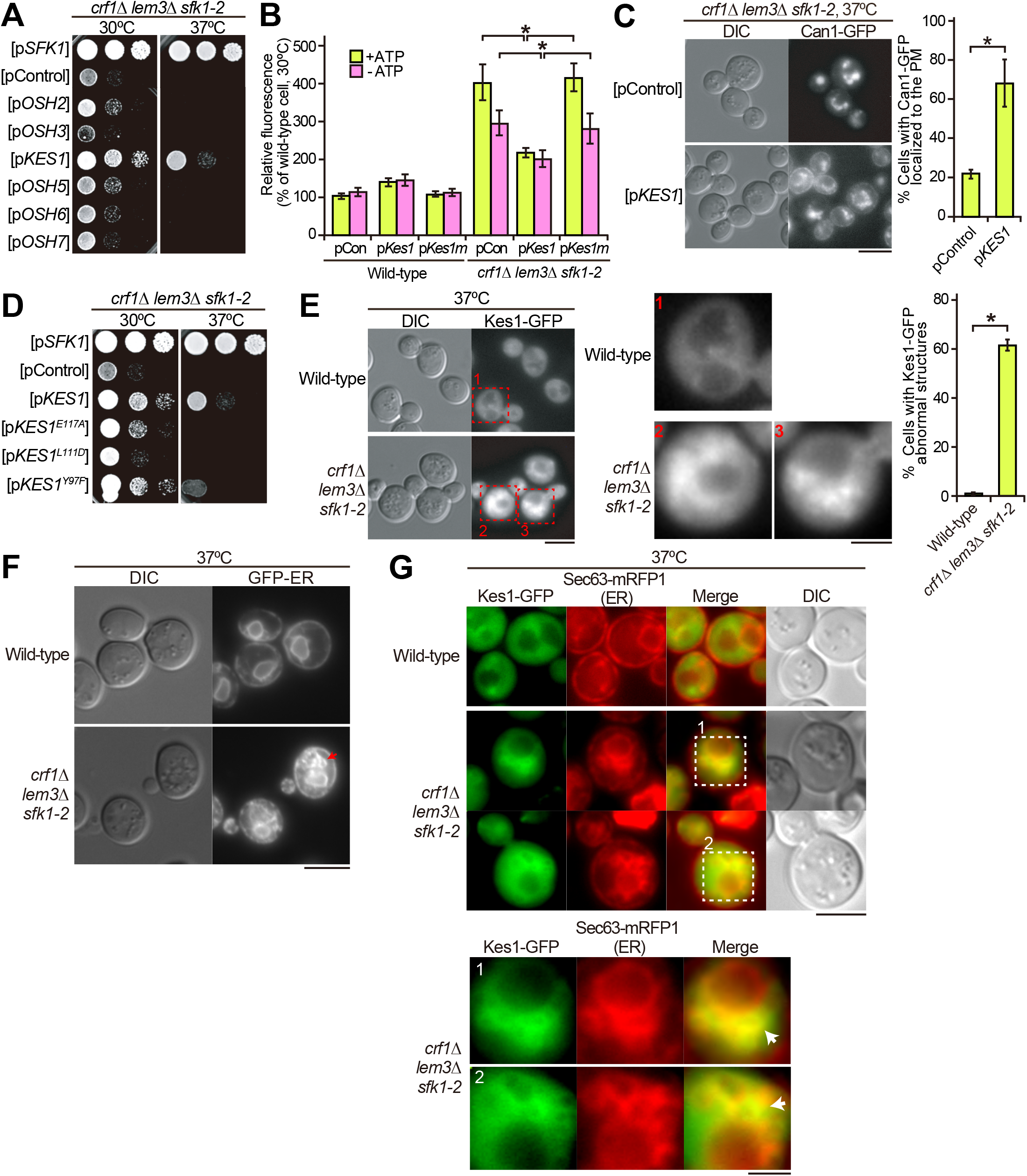
Overexpression of *KES1* partially suppresses the phenotypes in the *crf1*Δ *lem3*Δ *sfk1-2* triple mutant. (A) Suppression of the growth defect. Cell growth was examined in the *crf1*Δ *lem3*Δ *sfk1-2* triple mutant carrying pRS316-*SFK1*, YEplac195, or YEplac195-*OSH2-7*. YEplac195 is a multicopy plasmid. After cells were cultured in SDA-U medium at 30°C overnight, tenfold serial dilutions were spotted onto a YPDA plate, followed by incubation for 2 d at 30 or 37°C. (B) Suppression of high membrane permeability. Rhodamine uptake was examined in the wild-type and *crf1*Δ *lem3*Δ *sfk1-2* triple mutant harboring YEplac195 (pCon), YEplac195-*KES1* (p*Kes1*), or YEplac195-*KES1^L111D^* (p*Kes1m*). Cells were cultured as in Fig 2C, except that SDA-Ura medium was used. The rhodamine uptake assay was performed as described in the “Materials and Methods”. Rhodamine uptake is represented as a relative value of that (100%) in the wild-type harboring YEplac195 incubated at 30°C. Values represent the mean ± S.D. from three independent experiments. Asterisks indicate a significant difference, as determined by the Tukey–Kramer test (p < 0.05). (C) Suppression of the mislocalization of Can1-GFP. *CAN1-GFP*-expressing cells harboring YEplac195-*KES1-KanMX6* or YEplac195-*KanMX6* are shown. Cells were cultured as in Fig 2C, except that YPDA medium containing G418 was used. Bar, 5 µm. *Right panel*: the percentage of cells with Can1-GFP at the PM was determined and is expressed as the mean ± S.D. of three independent experiments (n > 263 cells in total for each strain). An asterisk indicates a significant difference, as determined by the Tukey–Kramer test (p < 0.05). (D) Overexpression of ergosterol-binding deficient *KES1* mutants does not suppress ts growth of the *crf1*Δ *lem3*Δ *sfk1-2* triple mutant. Cell growth was examined in the triple mutant transformed with pRS316-*SFK1*, YEplac195, YEplac195-*KES1*, or YEplac195-*KES1* mutants (E117A, L111D, and Y97F). Cells were spotted and grown as in (A). (E) Kes1-GFP is localized to abnormal structures in the *crf1*Δ *lem3*Δ *sfk1-2* triple mutant. *Left panel*: cells were cultured as in Fig 2C. Bar, 5 µm. *Middle panel*: enlarged images of dotted red squares in the left panel. Bar, 2 μm. *Right panel*: the percentage of cells harboring abnormal structures of Kes1-GFP was determined and is expressed as the mean ± S.D. of three independent experiments (n > 210 cells in total for each strain). An asterisk indicates a significant difference, as determined by the Tukey–Kramer test (p < 0.05). (F) Abnormal ER structures in the *crf1*Δ *lem3*Δ *sfk1-2* triple mutant. The ER was visualized by the expression of GFPenvy-Scs2^220–244^ (GFP-ER). Cells were cultured as in Fig 2C. An arrow indicates abnormal ER structures. Bar, 5 µm. (G) Colocalization of Kes1-GFP with Sec63-mRFP1 in the *crf1*Δ *lem3*Δ *sfk1-2* triple mutant. Cells were cultured as in Fig 2C. The abnormal ER structures in squares 1 and 2 are enlarged in the lower panel. Arrows represent Kes1-GFP puncta colocalized with Sec63-mRFP1. Bars, 2 µm (upper panel) and 0.4 μm (lower panel).

Kes1 interacts with phosphatidylinositol (PI) -4-phosphate (PI(4)P), and this interaction is implicated in sterol transport [67]. However, Sfk1 was implicated in the function of a PI 4 kinase, Stt4 [70, 71]. We thus examined the distribution of PI(4)P by using the PI(4)P-specific biosensor Osh2-PH in the *crf1*Δ *lem3*Δ *sfk1-2* triple mutant. However, the localization of Osh2-PH-GFP was not significantly affected (S5A Fig). The mammalian homolog of Sfk1, TMEM150A, interacts with PI 4 kinase type IIIα via its C-terminal domain [40, 71]. However, the C-terminally truncated *SFK1*Δ*C-GFP* did not show synthetic growth defects with *crf1*Δ *lem3*Δ mutations (S5B Fig). These results suggest that the defects in the *crf1*Δ *lem3*Δ *sfk1* triple mutants may not be closely related to PI(4)P.

We next examined whether the sterol-binding activity of Kes1 is required to suppress the growth defect of the *crf1*Δ *lem3*Δ *sfk1-2* triple mutant. Overexpression of *KES1*, *KES1^L111D^*, and *KES1^Y97F^*, which abolishes the binding of Kes1 to sterols [72], did not suppress the growth defect of the triple mutant (Fig 4D). Correspondingly, overexpression of *KES1^L111D^* did not suppress rhodamine accumulation (Fig 4B, p*Kes1m*). These results suggest that the *crf1*Δ *lem3*Δ *sfk1-2* triple mutant may have a defect in intracellular sterol transport.

We next examined the subcellular localization of Kes1 in the *crf1*Δ *lem3*Δ *sfk1-2* triple mutant (Fig 4E). Wild-type and double mutants displayed the diffuse distribution of Kes1-GFP in the cytosol with a few puncta (Fig 4E, S6A Fig). In contrast, the *crf1*Δ *lem3*Δ *sfk1-2* triple mutant showed Kes1-GFP localization in abnormal membranous structures (Fig 4E). Because they appeared to be localized around the nucleus, we examined the localization of the ER marker GFP-ER (GFPenvy-Scs2^220–244^) [73]. Abnormal ER structures close to the perinuclear ER were observed specifically in the triple mutant at a frequency similar to that of the abnormal Kes1-GFP structures (56%, n=210 cells) (Fig 4F, S6B Fig). Coexpression of *KES1-GFP* and the ER marker *SEC63-mRFP1* (monomeric red fluorescent protein 1 [*mRFP1*]) demonstrated that Kes1-GFP colocalized with Sec63-mRFP1 in the abnormal structures but not in the perinuclear ER in the *crf1*Δ *lem3*Δ *sfk1-2* triple mutant (Fig 4G). This colocalization was observed in 90% of the *crf1*Δ *lem3*Δ *sfk1-2* triple mutant displaying abnormal Kes1-GFP structures (n=450 cells). Taken together, these results suggest that the *crf1*Δ *lem3*Δ *sfk1-2* triple mutant may have a defect in ergosterol homeostasis.

### Ergosterol is reduced in the PM of the *crf1*Δ *lem3*Δ *sfk1* triple mutants

We examined the distribution of ergosterol in the *crf1*Δ *lem3*Δ *sfk1* triple mutants. Filipin is a polyene antibiotic that binds to cholesterol and ergosterol and is used as a probe for cellular sterol distribution [74, 75]. Wild-type and double mutants were evenly labeled with filipin at the PM (Fig 5A, S7 Fig). However, the *crf1 lem3*Δ *sfk1*Δ triple mutant drastically decreased filipin labeling to the PM and instead showed the enhancement of intracellular labeling (Fig 5A). This labeling pattern was similar to that of Kes1-GFP, but costaining experiments could not be performed because the fixation step for filipin staining diminished the GFP fluorescence. Eighty-three percent of the *crf1*Δ *lem3*Δ *sfk1*Δ triple mutant cells clearly displayed the loss or reduction of filipin signal in the PM (n=98 cells). Quantitative analysis of fluorescence images further confirmed the decrease in filipin intensity on the PM in the triple mutant (Fig 5A).

**Fig 5.**
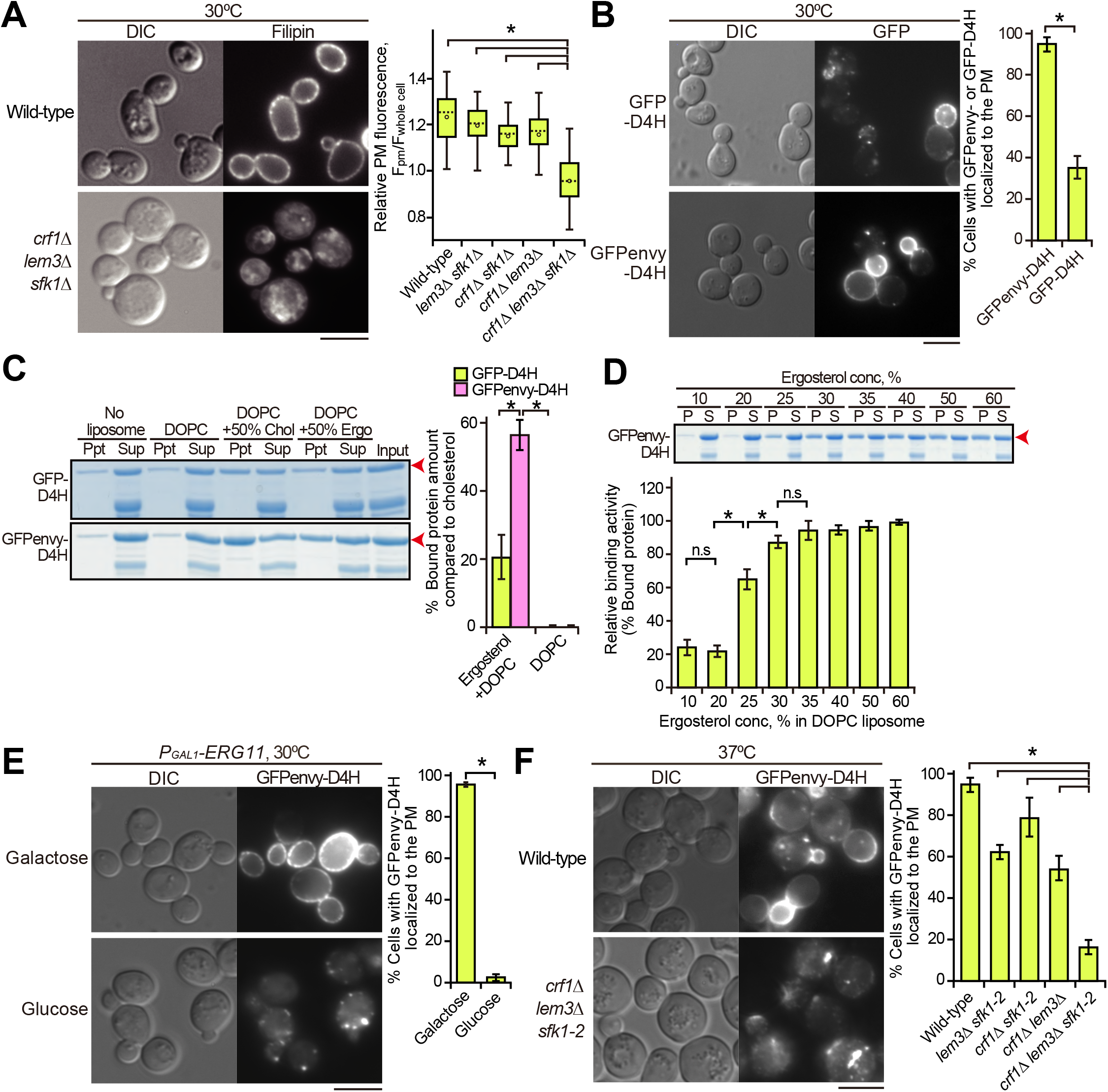
Ergosterol is decreased in the PM of the *crf1*Δ *lem3*Δ *sfk1* triple mutants. (A) Staining of ergosterol with filipin in the *crf1*Δ *lem3*Δ *sfk1*Δ triple mutant. Cells were grown in YPDA at 30°C, and filipin staining was performed as described in the “Materials and Methods”. *Right panel*: fluorescence intensity at the PM was quantitated as described in the “Materials and Methods”. The ratio of the fluorescence at the PM (F_pm_)/that of whole cell (F_whole cell_) was determined and expressed with a boxplot (whiskers: maximum and minimum values; box: first quartile, median, and third quartile; circle: average). The numbers of cells analyzed were 23, 38, and 35 for the wild-type, *lem3*Δ *sfk1*Δ, and *crf1*Δ *lem3*Δ *sfk1*Δ, respectively. An asterisk indicates a significant difference, as determined by the Tukey–Kramer test (p < 0.05). (B) The distribution of GFP- and GFPenvy-D4H in wild-type cells. Cells harboring pRS316-GFP-D4H or -GFPenvy-D4H were grown in SDA-Ura medium at 30°C to the mid-log phase. *Right panel*: the percentage of cells with GFP- or GFPenvy-D4H at the PM was determined and is expressed as the mean ± S.D. of three independent experiments (n > 155 cells in total for each strain). An asterisk indicates a significant difference, as determined by the Tukey–Kramer test (p < 0.05). (C) GFPenvy-D4H binds to ergosterol *in vitro*. The liposome sedimentation assay was performed as described in the “Materials and Methods”. Proteins were incubated with liposomes composed of DOPC or DOPC and 50% (mol) ergosterol (Ergo) or cholesterol (Chol) (with corresponding reduction in DOPC), followed by pelleting by centrifugation. Supernatant (Sup) and pellet (Ppt) fractions were separated by SDS-PAGE, followed by staining with Coomassie Brilliant Blue. Arrowheads indicate GFP-D4H or GFPenvy-D4H. A lower band appears to be an incomplete fragment. *Right panel*: proteins bound to liposomes were quantitated by an image analyzer. The percentage of the bound protein was determined as a relative value of that bound to cholesterol liposomes and is expressed as the mean ± S.D. of three independent experiments. Asterisks indicate a significant difference, as determined by the Tukey–Kramer test (p < 0.05). (D) GFPenvy-D4H binds to DOPC liposomes containing more than 25% ergosterol. A liposome sedimentation assay was performed as in (C) with DOPC liposomes containing the indicated mol % ergosterol. An arrowhead indicates GFPenvy-D4H. A lower band appears to be an incomplete fragment. The amount of protein bound to liposomes is expressed as a relative value (percentage) of that bound to 60% ergosterol liposomes. Values represent the mean ± S.D. of three independent experiments. “S” and “P” indicate supernatant and pellet fractions, respectively. Asterisks and “n.s” indicate significant and no significant differences as determined by the Tukey–Kramer test (*p < 0.05), respectively. (E) The PM localization of GFPenvy-D4H was dependent on ergosterol. *ERG11* was expressed under the control of the glucose-repressible *P_GAL1_* promoter. Cells were grown in SGA-Ura medium to the mid-log phase at 30°C and then inoculated into fresh galactose (SGA-Ura) or glucose (SDA-Ura) medium, followed by incubation for 12 h at 30°C. *Right panel*: the percentage of cells with GFPenvy-D4H at the PM was determined and is expressed as the mean ± S.D. of three independent experiments (n > 108 cells in total for each condition). An asterisk indicates a significant difference, as determined by the Tukey–Kramer test (p < 0.05). (F) GFPenvy-D4H was not localized to the PM in the *crf1*Δ *lem3*Δ *sfk1-2* triple mutant. Cells were cultured as in Fig 2C, except that SDA-Ura medium was used. *Right panel*: the percentage of cells with GFPenvy-D4H at the PM was determined and is expressed as the mean ± S.D. of three independent experiments (n > 253 cells in total for each strain). An asterisk indicates a significant difference, as determined by the Tukey–Kramer test (p < 0.05). Bars, 5 µm.

We also examined another sterol biosensor, D4H. A bacterial protein toxin, perfringolysin O, binds to cholesterol via its domain 4 (D4) [76, 77]. A D4 derivative, D4H (D4^D434S^), has been developed as a more sensitive probe; it binds to liposomes containing 20-30% cholesterol mole concentration [48, 77]. Although D4H has recently been used to detect the PM sterol in fission yeast [78], D4H has not been applied to budding yeast. We generated two fluorescent protein-conjugated D4Hs, GFP-D4H and GFPenvy-D4H, in which GFPenvy is a photostable dimeric GFP derivative [79, 80]. When expressed in wild-type cells, GFP-D4H was localized to the PM in 35% of the cells (Fig 5B). In contrast, GFPenvy-D4H was localized to the PM in 94% of the cells. Interestingly, GFPenvy-D4H showed a characteristic localization pattern; it preferentially localized to daughter cells compared to mother cells. This localization pattern is described in the last part of “Results” in more detail.

We next examined the binding activity of recombinant GFP-D4H and GFPenvy-D4H to ergosterol *in vitro* by liposome sedimentation assay using 1,2-Dioleoyl-sn-glycero-3-phosphocholine (DOPC) liposomes containing either 50% cholesterol or 50% ergosterol. Consistent with a previous report [81], GFP-D4H bound to ergosterol liposomes at an efficiency of 20% of that to cholesterol liposomes (Fig 5C). On the other hand, GFPenvy-D4H bound to ergosterol liposomes at an efficiency of 56% of that to cholesterol liposomes (Fig 5C), consistent with the results in living cells. The affinity of GFPenvy-D4H to ergosterol was examined with DOPC liposomes containing different concentrations of ergosterol from 10 to 60 mol %. Binding was detected when the ergosterol concentration was 25% or higher (Fig 5D). These results are comparable to the affinity of D4H to cholesterol [48]. The higher affinity of GFPenvy-D4H to ergosterol than GFP-D4H might be because GFPenvy forms a dimeric structure [80].

We next confirmed that GFPenvy-D4H binds to ergosterol in living cells. The *ERG11* gene encodes lanosterol demethylase, which is essential for ergosterol synthesis [82]. Both the shut-off of *ERG11* gene expression and treatment with the Erg11 inhibitor fluconazole inhibited GFPenvy-D4H distribution to the PM (Fig 5E, S8A Fig). We also examined the localization of GFPenvy-D4H in mutants of genes involved in the late steps of the ergosterol biosynthesis pathway (*ERG2-6*) [83, 84]. GFPenvy-D4H was not localized to the PM except for *erg4*Δ, which catalyzes the last step (S8B Fig). These results suggest that GFPenvy-D4H is localized to the PM by binding to ergosterol.

We examined the distribution of GFPenvy-D4H in the *crf1*Δ *lem3*Δ *sfk1-2* triple mutant. The localization of GFPenvy-D4H to the PM decreased to some extent in the *lem3*Δ *sfk1-2* and *crf1*Δ *lem3*Δ double mutants, but it drastically decreased to 16% in the *crf1*Δ *lem3*Δ *sfk1-2* triple mutant (Fig 5F, S8C Fig). Together with the results of filipin staining, we concluded that ergosterol is significantly lost from the PM in the *crf1*Δ *lem3*Δ *sfk1* triple mutants. Kes1 overexpression increased the PM localization of GFPenvy-D4H from 25% to 55% in the *crf1*Δ *lem3*Δ *sfk1-2* triple mutant (S8D Fig). These results suggest that the increased Kes1 enhances ergosterol transport to or inhibits loss of ergosterol from the PM in the triple mutant and that loss of ergosterol from the PM causes phenotypes of the triple mutant, including the growth defect.

We next examined whether exogenously added ergosterol would suppress the growth defect in the *crf1*Δ *lem3*Δ *sfk1-2* triple mutant. We used strains carrying the gain-of-function mutation of a transcription factor *UPC2*, *upc2-1* (G888D), which results in increased uptake of exogenous ergosterol under aerobic conditions [85, 86]. The exogenously added ergosterol, which was sufficient to suppress the growth defect of the ergosterol-deficient *hem1*Δ mutant [87], did not suppress the growth defect of the *crf1*Δ *lem3*Δ *sfk1-2* triple mutant even when *KES1* was overexpressed (Fig 6A). We presumed that the triple mutant cannot retain exogenously added ergosterol in the PM. Thus, we monitored the distribution of exogenously added TopFluor-cholesterol (TF-Chol), a fluorescent dye-conjugated cholesterol analog [88]. TF-Chol was retained in the PM of the wild-type and double mutants containing *upc2-1* but was not in the *crf1*Δ *lem3*Δ *sfk1-2 upc2-1* mutant; only 23% of the triple mutant showed TF-Chol in the PM (Fig 6B, S9 Fig). TF-Chol appeared to be internalized into the cell to be incorporated into cytoplasmic punctate structures in the *crf1*Δ *lem3*Δ *sfk1-2 upc2-1* mutant (Fig 6B). These results suggest that ergosterol is not retained in the PM and that it is transported to intracellular punctate structures in the *crf1*Δ *lem3*Δ *sfk1-2* mutant.

**Fig 6.**
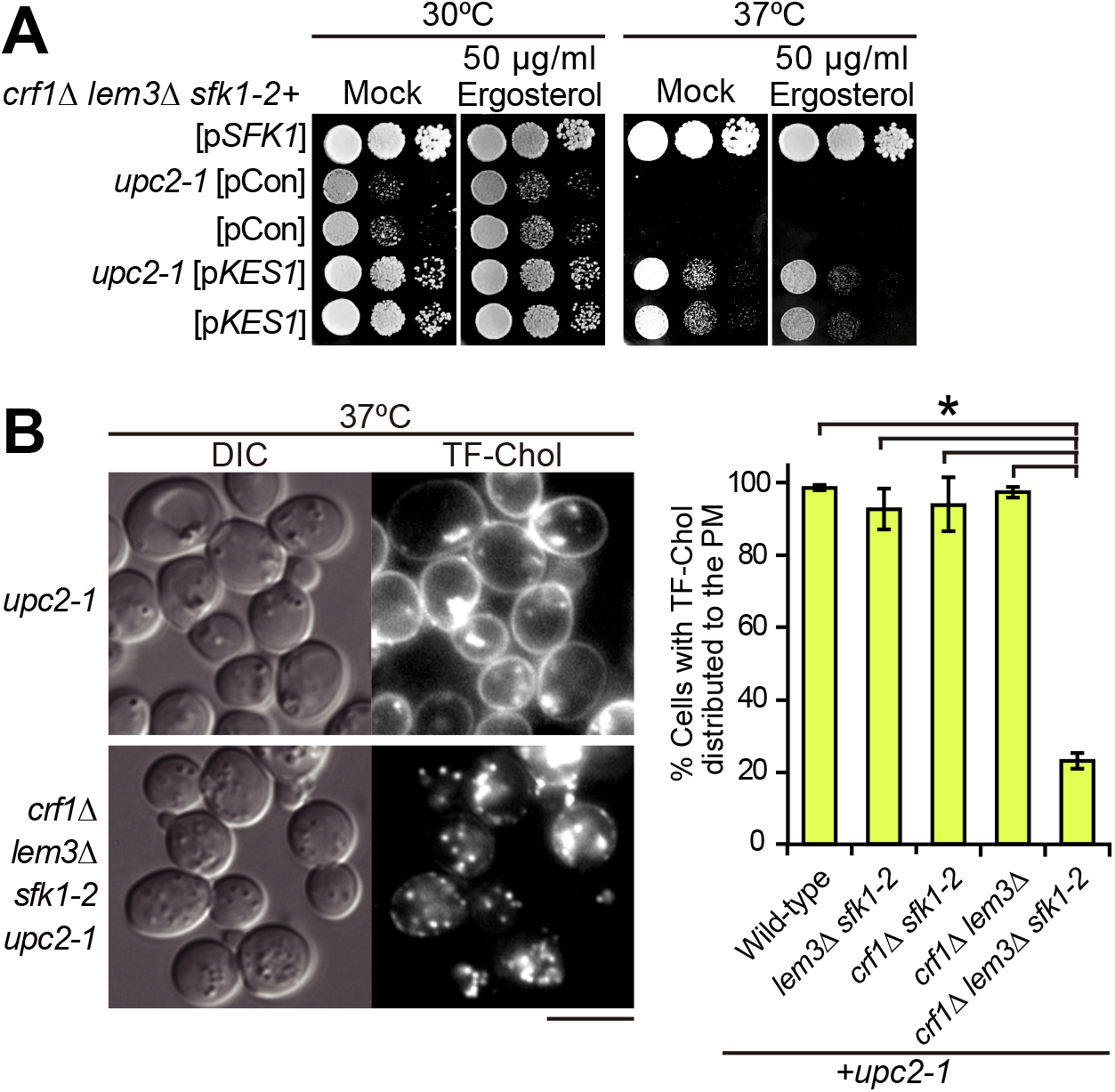
Exogenously added ergosterol appears to not be retained in the PM of the *crf1*Δ *lem3*Δ *sfk1-2 upc2-1* mutant. (A) Exogenous ergosterol did not suppress ts growth in the *crf1*Δ *lem3*Δ *sfk1-2 upc2-1* mutant. The *crf1*Δ *lem3*Δ *sfk1-2* mutant with or without the *upc2-1* mutation was transformed with pRS315-*SFK1*, YEplac181, or YEplac181-*KES1*. After cells were cultured in SD-Ura-Leu medium at 30°C overnight, tenfold serial dilutions were spotted onto a YPDA plate containing 0.5% Tween-80 and 0.5% ethanol with or without 50 µg/mL ergosterol, followed by incubation for 1.5 d at 30 or 37°C. (B) TF-Chol is not retained in the PM of the *crf1*Δ *lem3*Δ *sfk1-2 upc2-1* mutant. *Left panel*: cells were cultured and labeled with TF-Chol as described in the “Materials and Methods”. Bar, 5 µm. *Right panel*: the percentage of cells with TF-Chol at the PM was determined and is expressed as the mean ± S.D. of three independent experiments (n > 290 cells in total for each strain). An asterisk indicates a significant difference, as determined by the Tukey–Kramer test (p < 0.05).

### Ergosterol is esterified and accumulated in LDs in the *crf1*Δ *lem3*Δ *sfk1-2* mutant

The loss of ergosterol in the PM raises a question: where does ergosterol distribute in the cell? We performed thin-layer chromatography (TLC) analysis of total sterols extracted from the cells. In double mutants, the free ergosterol level was approximately 80-85% of that in the wild-type, but it decreased to 50% in the *crf1*Δ *lem3*Δ *sfk1*Δ triple mutant (Fig 7A). TLC analysis also showed a large increase in esterified ergosterol in the *crf1*Δ *lem3*Δ *sfk1*Δ triple mutant. We confirmed that this spot was observed in the wild-type at the stationary phase, but not in the acyl-CoA:sterol acyltransferase deficient *are1 are2* mutant (S10 Fig) [89]. Because esterified ergosterol is the main component of LDs, these results suggest that LDs are increased in the *crf1*Δ *lem3*Δ *sfk1*Δ triple mutant. To confirm this, we stained LDs with the lipophilic dye Nile red, which stains neutral lipids in LDs, triacylglycerol and esterified ergosterol [90]. Neither the wild-type nor the double mutants showed obvious staining of Nile red, whereas the *crf1*Δ *lem3*Δ *sfk1*Δ triple mutant exhibited a clear increase in cells showing Nile red puncta (Fig 7B, S11A Fig). We further examined the localization of GFP-tagged LD-related proteins, Tgl1 (steryl ester lipase) [91] and Faa4 (long-chain fatty-acid-CoA ligase) [92]. The wild-type and the double mutants contained a few puncta of these proteins, whereas the numbers of Tgl1-GFP and Faa4-GFP puncta increased 2.5- and 2.8-folds, respectively, in the *crf1*Δ *lem3*Δ *sfk1-2* triple mutant compared to those in the wild-type (Fig 7C, S11B Fig). We confirmed that Tgl1-GFP and Faa4-GFP puncta were colocalized with Nile red-positive puncta; 85% of Tgl1-GFP (n=377) and 88% of Faa4-GFP (n=453) puncta were colocalized with Nile red in the *crf1*Δ *lem3*Δ *sfk1-2* triple mutant (Fig 7C). These results suggest that a substantial amount of ergosterol was esterified and accumulated in LDs in the *crf1*Δ *lem3*Δ *sfk1* triple mutants.

**Fig 7.**
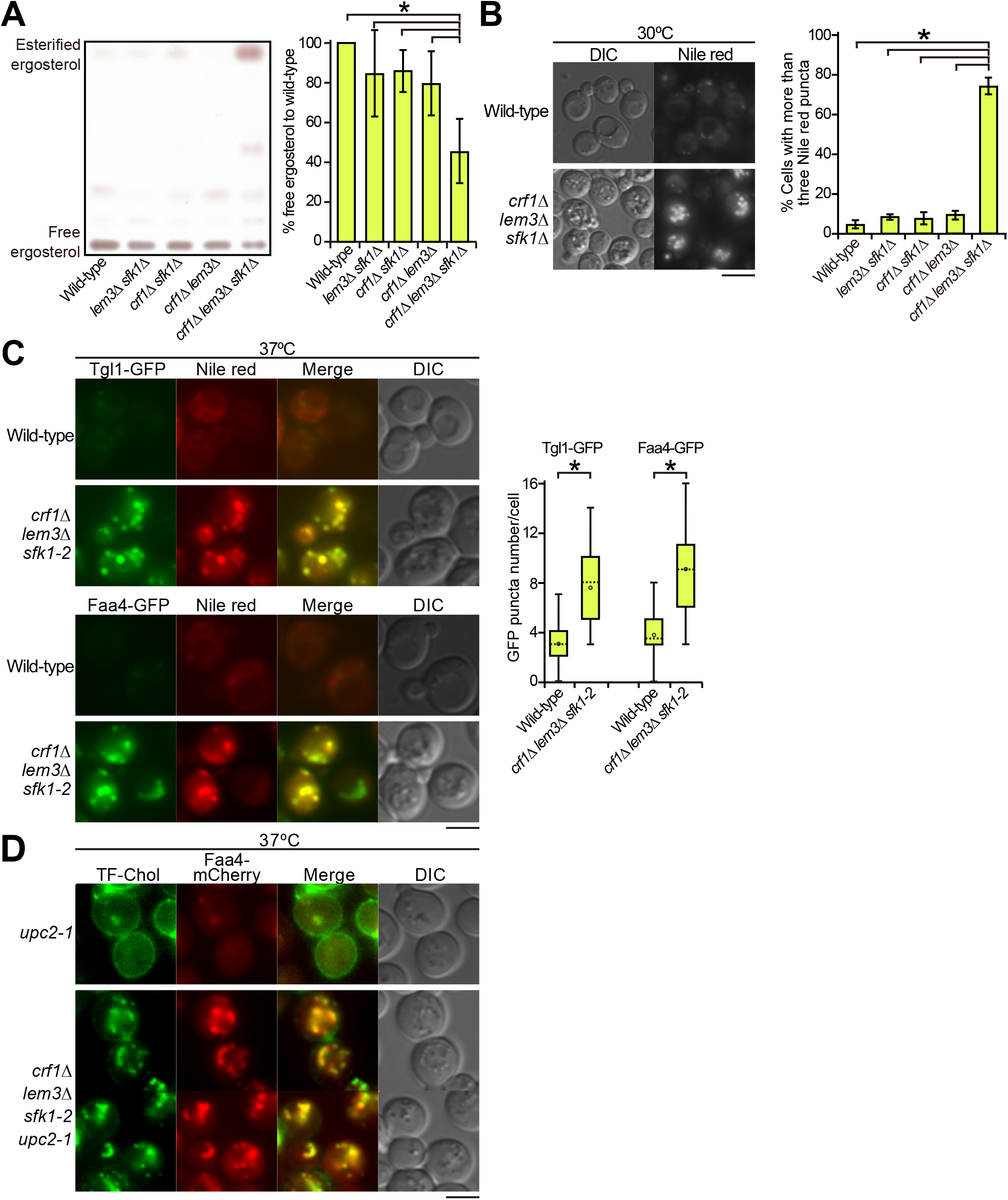
LDs were increased in the *crf1*Δ *lem3*Δ *sfk1* triple mutants. (A) TLC analysis of total sterols. Ergosterol contents were analyzed by TLC as described in the “Materials and Methods”. *Right panel*: the percentage of free ergosterol relative to that of the wild-type was determined and is expressed as the mean ± S.D. derived from the analysis of six independent samples. An asterisk indicates a significant difference, as determined by the Tukey–Kramer test (p < 0.05). (B) Increase in Nile red-positive puncta in the *crf1*Δ *lem3*Δ *sfk1*Δ triple mutant. Cells were cultured in YPDA medium to the mid-log phase at 30°C, followed by Nile red staining. Nile red staining was performed as described in the “Materials and Methods”. *Right panel*: the percentage of cells with more than three Nile Red puncta was determined and is expressed as the mean ± S.D. of three independent experiments (n > 315 cells in total for each strain). An asterisk indicates a significant difference, as determined by the Tukey–Kramer test (p < 0.05). (C) Increase in LD marker (Tgl1-GFP and Faa4-GFP)-containing structures in the *crf1*Δ *lem3*Δ *sfk1-2* triple mutant. Cells were cultured as in Fig 2C, followed by Nile red staining. *Right panel*: the numbers of Tgl1-GFP or Faa4-GFP puncta were counted in a single focal plane of each cell and expressed with boxplots (whiskers: maximum and minimum values; box: first quartile, median, and third quartile; circle: average). The number of cells analyzed was 51 and 50 (wild-type) and 53 and 51 (triple mutant) for Tgl1-GFP and Faa4-GFP, respectively. Asterisks indicate a significant difference, as determined by the Tukey–Kramer test (p < 0.05). (D) Colocalization of TF-Chol puncta with Faa4-mCherry in the *crf1*Δ *lem3*Δ *sfk1-2 upc2-1* mutant. Cells were cultured as in Fig 2C, except that YPDA medium containing TF-Chol was used. Bars, 5 µm.

TF-Chol was also detected in intracellular puncta in the *crf1*Δ *lem3*Δ *sfk1-2 upc2-1* mutant (Fig 6B). We next examined whether TF-Chol colocalizes with Faa4-mCherry in the triple mutant. The *crf1*Δ *lem3*Δ *sfk1-2 upc2-1* mutant contained approximately 9∼14 TF-Chol puncta per cell, and 88% of these puncta (n=603) were colocalized with Faa4-mCherry (Fig 7D).

Taken together, these results suggest that ergosterol is not retained in the PM and is transported to LDs in an esterified form, probably via the ER, in the *crf1*Δ *lem3*Δ *sfk1* triple mutants.

### The inhibition of sterol esterification partially suppresses growth defects and sterol retention in the PM of the *crf1***Δ** *lem3***Δ** *sfk1-2* mutant

We examined whether inhibition of sterol esterification by mutations in *ARE1*/*ARE2* suppresses the phenotypes of the *crf1*Δ *lem3*Δ *sfk1-2* triple mutant. The growth defect of the triple mutant was partially suppressed by the *are2*Δ mutation but not by the *are1*Δ mutation (Fig 8A). Consistently, Are2 accounts for 65–75% of total cellular acyl-CoA:sterol acyltransferase activity [93, 94]. The growth defect of the triple mutant was not suppressed by either the *dga1*Δ or *lro1*Δ mutation, which abolishes the synthesis of triacylglycerol [95–97], the other major lipid in LDs (S12A Fig). We then examined whether the *are2*Δ mutation restored sterol retention in the PM in the *crf1*Δ *lem3*Δ *sfk1-2* triple mutant. The *are2*Δ mutation increased the number of cells showing the PM localization of GFPenvy-D4H from 17% to 60% in the *crf1*Δ *lem3*Δ *sfk1-2* mutant (Fig 8B). A TF-Chol uptake experiment could not be performed because the *are2*Δ *upc2-1* double mutant showed a severe growth defect (S12B Fig). These results are consistent with our notion that loss of ergosterol from the PM is responsible for the growth defect of the *crf1*Δ *lem3*Δ *sfk1-2* triple mutant.

**Fig 8.**
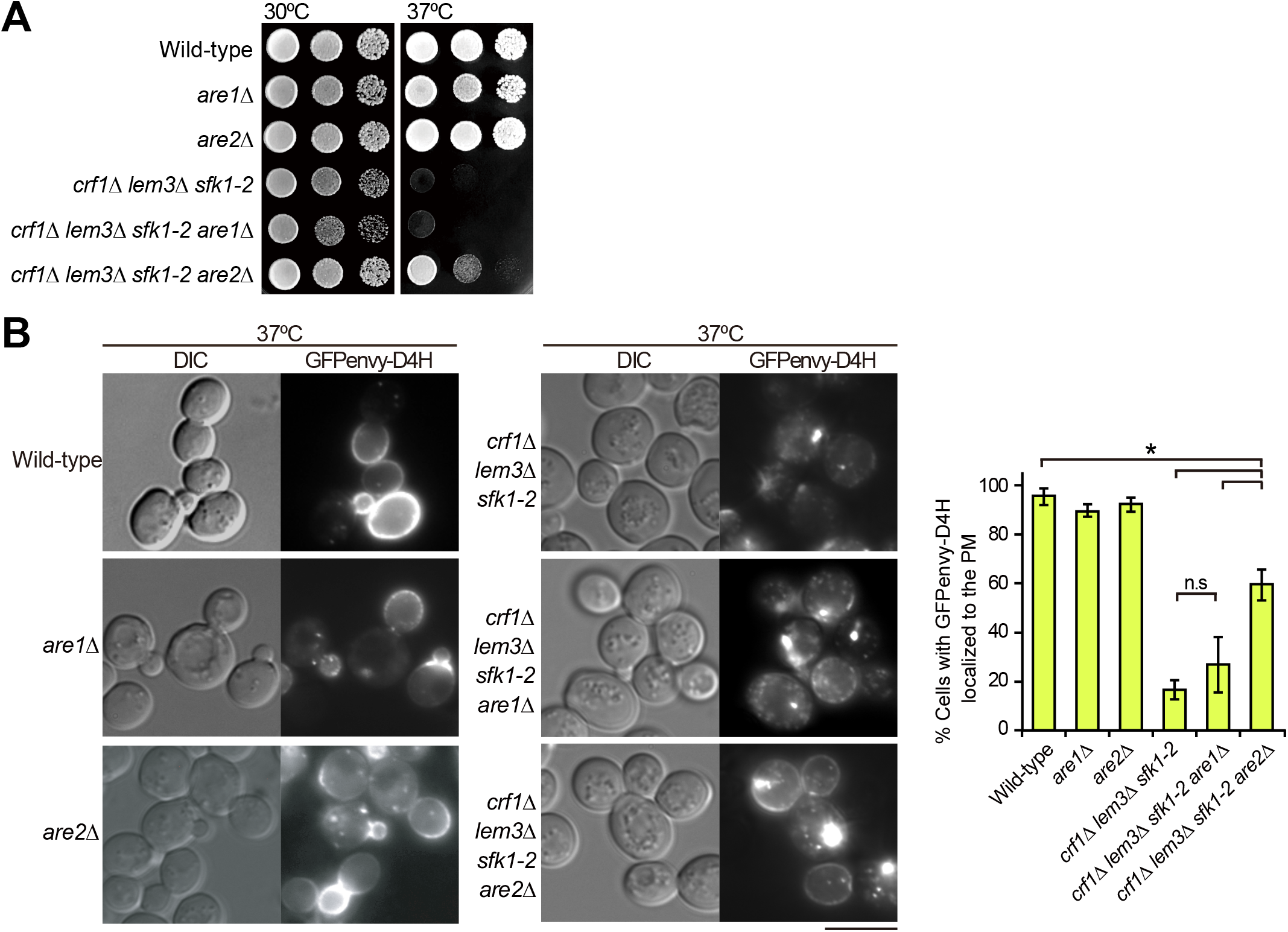
The *are2*Δ mutation partially restores ergosterol in the PM of the *crf1*Δ *lem3Δ sfk1-2* triple mutant. (A) Suppression of the growth defect. Tenfold serial dilutions were spotted onto a YPDA plate, followed by incubation for 1.5 d at 30 or 37°C. (B) Restoration of GFPenvy-D4H localization to the PM. Cells were cultured as in Fig 2C except that SDA-Ura medium was used. Bar, 5 µm. *Right panel*: the percentage of cells with GFPenvy-D4H at the PM was determined and is expressed as the mean ± S.D. of three independent experiments (n > 219 cells in total for each strain). Asterisks and “n.s” indicate significant and no significant differences as determined by the Tukey–Kramer test (*p < 0.05), respectively.

### Overexpression of Sfk1 alters the localization of GFPenvy-D4H

The molecular function of Sfk1 remains to be clarified, but our results described above may suggest that Sfk1 is functionally related to ergosterol in the PM. We examined whether a mutation or overexpression of *SFK1* affects the localization of GFPenvy-D4H.

GFPenvy-D4H exhibited polarized localizations in many wild-type cells; it was localized to daughter cells (buds) or near the bud neck in medium- or large-budded cells (Fig 9A, yellow and pink arrows). These results suggest that the accessibility to ergosterol is different between bud and mother PMs because filipin, which was used in fixed cells, evenly stained ergosterol in daughter and mother cells. Interestingly, Sfk1 was mainly localized to mother cells but not to daughter cells, as described previously [71], showing a localization pattern opposite that of GFPenvy-D4H (Fig 9B and 9C). The GFPenvy-D4H localizations in large-budded cells were categorized into three patterns: (1) localized throughout the PM (not polarized), (2) localized to the bud and mother cell PM near the bud neck (partially polarized), and (3) localized only to the bud (polarized). These differences may be because GFPenvy-D4H was expressed from a centromeric plasmid. The fluorescence intensity profiles of Sfk1-3xmCherry and GFPenvy-D4H are shown for the “polarized” (Fig 9C) and “partially polarized” (S13A Fig) patterns. The proportion of these localization patterns was not changed in the *sfk1*Δ mutant (Fig 9A and 9D).

**Fig 9.**
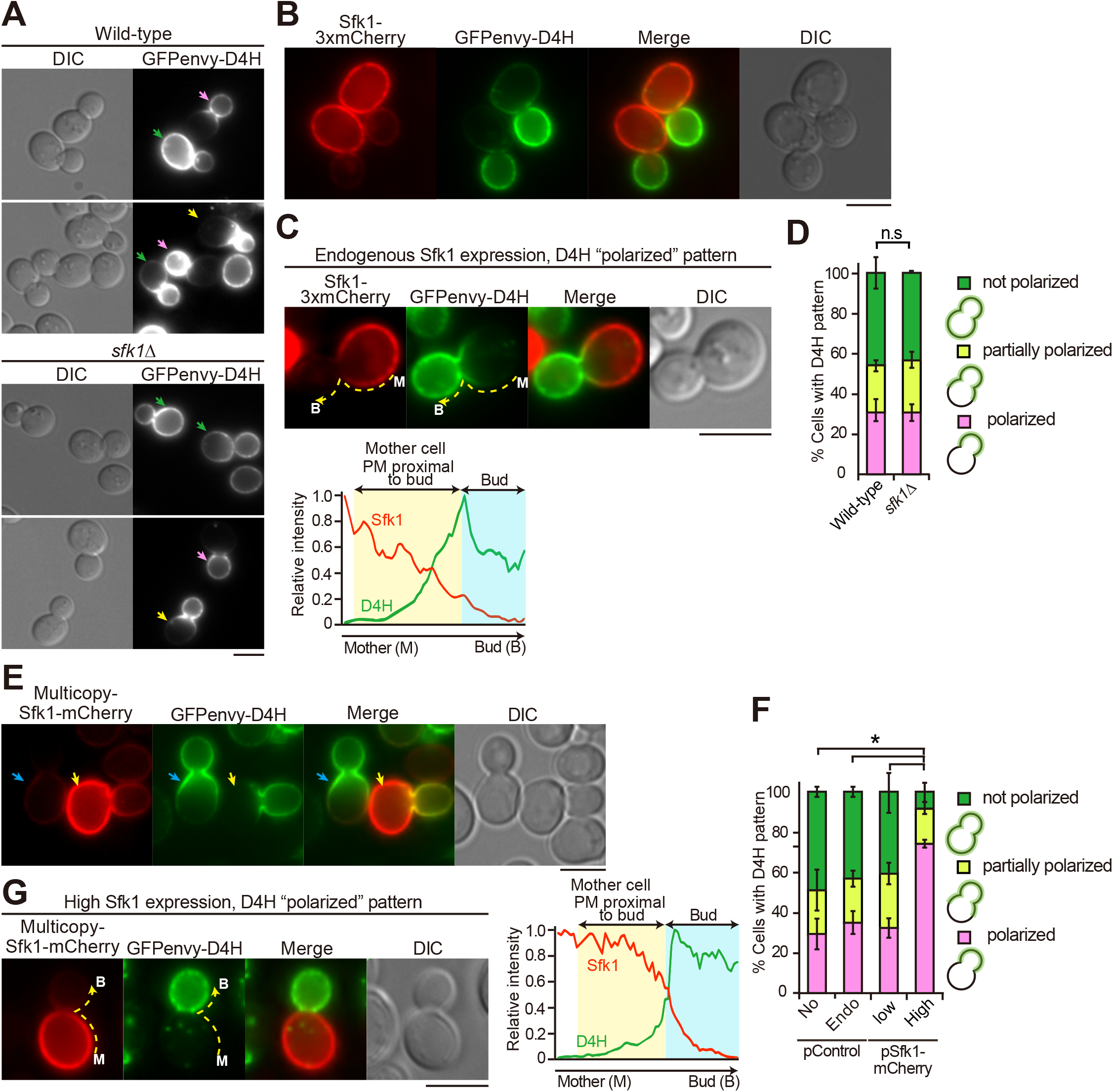
Overexpression of Sfk1 excludes GFPenvy-D4H from the mother cell PM. (A) The polarized distribution of GFPenvy-D4H. Wild-type or *sfk1*Δ cells carrying pRS316-GFPenvy-D4H were grown in SD-Ura medium to the mid-log phase at 30°C. “Polarized”, “partially polarized”, and “not polarized” localizations of GFPenvy-D4H are indicated with pink, yellow, and green arrows, respectively. (B) Complementary localization of GFPenvy-D4H and Sfk1-3xmCherry to daughter (bud) and mother cells, respectively. Wild-type cells expressing these proteins were grown at 30°C. To show endogenously expressed Sfk1-3xmCherry clearly, the brightness was adjusted to make it brighter. (C) Fluorescence intensity profile of a cell showing the “polarized” pattern of GFPenvy-D4H. Fluorescence signals were quantified along the dotted line from the mother cell to the bud. The brightness of Sfk1-3xmCherry was adjusted as in (B). (D) Quantification of three GFPenvy-D4H localization patterns. The cells in (A) were examined. The percentage of cells showing “polarized”, “partially polarized”, and “not polarized” localizations of GFPenvy-D4H was determined as described in the “Materials and Methods” and is expressed as the mean ± S.D. of three independent experiments (n > 150 cells in total for each strain). “n.s,” indicates no significant difference between all combinations as determined by the Tukey–Kramer test. (E) Heterogeneous (high and low) expression of Sfk1-mCherry by a multicopy plasmid. Wild-type cells carrying pRS316-GFPenvy-D4H and YEplac181-*SFK1-mCherry* were grown in SD-Leu-Ura medium to the mid-log phase at 30°C. The brightness was not adjusted after background subtraction. Arrows indicate cells highly (yellow) and lowly (cyan) expressing Sfk1-mCherry. (F) High expression of Sfk1-mCherry significantly increased the “polarized” pattern of GFPenvy-D4H. Cells were examined and categorized as in (D). Low or high expression of Sfk1-mCherry was determined as described in the legend of S13B Fig. *Bars*: No, control plasmid; Endo, endogenous expression of Sfk1-3xmCherry; Low, multicopy plasmid of *SFK1-mCherry* but low expression of Sfk1-mCherry; High, multicopy plasmid of *SFK1-mCherry* and high expression of Sfk1-mCherry. The percentage of cells showing indicated patterns is expressed as the mean ± S.D. of three independent experiments (n > 103 cells in total for each strain). An asterisk indicates a significant difference, as determined by the Tukey–Kramer test (p < 0.05), in the “polarized” and “not polarized” patterns. (G) GFPenvy-D4H was exclusively distributed to the bud in a cell highly expressing Sfk1-mCherry. The brightness is not adjusted after background subtraction. The right panel represents the fluorescence intensity profile quantified as in (C). Bars, 3 µm.

We then examined the effect of *SFK1* overexpression by using a multicopy plasmid carrying *SFK1-mCherry*. Expression from a multicopy plasmid generates heterogeneity in the level of gene expression among individual cells because of variation in plasmid copy number [98]. We took advantage of this expression characteristic to examine the correlation between the Sfk1 expression level and the D4H localization pattern. Cells were categorized into high and low expression groups based on the fluorescent intensity of Sfk1-mCherry. The relative expression level of *SFK1-mCherry* in highly expressing cells was more than 3-fold that in lowly expressing cells (S13B Fig). In cells lowly expressing Sfk1-mCherry, the GFPenvy-D4H localization pattern was not changed (Fig 9E, cyan arrows, and 9F). The mother cell-specific localization pattern of Sfk1-mCherry was not changed in highly expressing cells (Fig 9E, yellow arrows). Interestingly, in cells highly expressing Sfk1-mCherry, GFPenvy-D4H distribution was restricted exclusively to the daughter cells, and those cells that showed the “polarized” pattern were largely increased to 74% (Fig 9F). In these cells, the fluorescence intensity of GFPenvy-D4H was weak in the mother cell PM but increased sharply near the bud neck (Fig 9G). These results suggest that Sfk1 might maintain ergosterol in a state that is inaccessible to GFPenvy-D4H, although its function may be redundant with that of an unknown protein.

## Discussion

More than two decades have passed since the first report on the asymmetric distribution of phospholipids in the PM [3], but our understanding of its physiological significance is still limited. Our genetic screening reveals that the loss of Dnf1/2-Lem3 and Dnf3-Crf1 flippases and Sfk1 results in severe growth defects, provably due to loss of ergosterol from the PM. Dnf3 was shown to be involved in some PM functions, including mating pheromone signaling [21] and pseudohyphal growth [35], but this is the first demonstration of Dnf3 involvement in essential cell function in a vegetative cell.

Disruption of phospholipid asymmetry is one main reason for the loss of ergosterol from the PM in the triple mutant. Phospholipids interact with sterol via their headgroups and acyl chains, which contributes to ordering membrane lipids and securing lipid packing [99]. In the *crf1*Δ *lem3*Δ *sfk1* triple mutants, PS and PE are more exposed to the extracellular leaflet than in the double mutants, and the level of PS in the cytoplasmic leaflet appears to be decreased. Sterols have a higher affinity to phospholipids containing saturated acyl chains than those containing unsaturated acyl chains [99], and PS and PE species in the PM are more abundant in those containing saturated acyl chains than in other organelles in budding yeast [100]. In addition, according to the umbrella model [99], phospholipid head groups in the membrane shield nonpolar cholesterol bodies from the aqueous phase. PS with a large headgroup has a higher affinity for cholesterol than other phospholipids [48, 101]. Therefore, ergosterol, which is also enriched in the cytoplasmic leaflet [102], loses favorable interacting partners in the triple mutant. This would result in a vast increase in “active ergosterol”, which may be actively removed from the PM by sterol transfer proteins (STPs) (see below).

Although Sfk1 is implicated in the regulation of phospholipid asymmetry (Fig 2) [39], its protein function remains unknown. Our results that overexpression of Sfk1 excludes GFPenvy-D4H from the mother cell PM suggest that Sfk1 may be functionally relevant to ergosterol. GFPenvy-D4H preferentially localized to the daughter cell PM. However, the ergosterol contents in the cytoplasmic leaflet did not seem to be significantly different between daughter and mother cell PMs because filipin uniformly stained both membranes and because ergosterol is predominantly localized to the cytoplasmic leaflet of the PM, including mother cells [102]. Thus, the differential GFPenvy-D4H localization might reflect different physical states of ergosterol. When sterol levels exceed the interacting capacity of phospholipids in the membrane, a sterol molecule is predicted to be exposed to the surface of the membrane, which increases the chance of its interaction with sterol-sensing proteins. The model defines this behavior as the chemical activation of cholesterol [99, 103–107]. It has been suggested that the D4-containing domain of perfringolysin O preferentially interacts with active cholesterol [104]. Thus, we propose that the chemical activity of ergosterol is higher in the PM of daughter cells than in that of mother cells. The PM of daughter cells is mainly made from newly synthesized lipids by polarized vesicle transport [108]. In this membrane, PE and PS are exposed, and Dnf1/2-Lem3 flippases actively flip these phospholipids to the cytoplasmic leaflet [12, 28]. Thus, phospholipid asymmetry seems to be more established in the PM of mother cells. Sfk1, which is mainly localized to the mother cell PM, might be involved in the maintenance of phospholipid asymmetry; our previous results suggest that Sfk1 represses spontaneous transbilayer movement of phospholipids [39]. This is consistent with the notion that Sfk1 may promote lipid packing in the PM. An interesting possibility is that Sfk1 enhances interactions between ergosterol and phospholipids, and thus, Sfk1 decreases active ergosterol.

The main localization sites were different between Dnf1/2-Lem3 and Sfk1, and the localization of Dnf3-Crf1 in the PM has not been clearly shown in the wild-type. However, simultaneous loss of these proteins leads to severe disorganization of the PM, in which active ergosterol would be highly increased due to a reduced shielding effect by phospholipids and reduced lipid packing. The ergosterol in the triple mutant appears to be highly accessible and easily extracted by STPs, resulting in the loss of ergosterol from the PM. Some STPs, including those yet to be identified, seem to be involved in sterol transfer from the PM. These include oxysterol binding protein homologs (Osh) and lipid transfer proteins anchored at a membrane contact site (LAMs) with StARkin domains [67, 109].

Our finding that ergosterol lost from the PM accumulated in LDs as esterified ergosterol is consistent with studies using exogenously added ergosterols [87, 110]. The PM has a much higher ergosterol concentration than the ER, but ergosterol transport by STPs between these membranes is kept in equilibrium because the active ergosterol concentration seems to be similar in these membranes; the ER membrane contains less saturated phospholipids, and thus ergosterol in the ER is not shielded by surrounding phospholipids [109]. In the *crf1*Δ *lem3*Δ *sfk1* triple mutants, a vast increase in active ergosterol occurs in the PM, and STPs transport these ergosterols to the ER, in which ergosterol is esterified by Are1/Are2 to form LDs, until the active ergosterol concentration in the PM is balanced with that in the ER. The *are2*Δ mutation increases active ergosterol in the ER due to defective esterification, and this ergosterol would be transferred to the PM, resulting in the partial suppression of growth defects in the triple mutant. Interestingly, in the triple mutant, abnormal ER structures accumulated around the perinuclear ER, and Kes1-GFP was localized to these structures. Because overexpression of Kes1 partially suppressed the growth defect of the triple mutant, those Kes1-GFP signals might reflect Kes1 in the transport cycle of ergosterol between the ER and the PM.

How sterols are maintained at a high concentration in the PM has been a longstanding question in membrane biology. Phospholipid asymmetry, a conserved feature in the PM, has been implicated in this role [6], but genetic analyses of flippases did not clearly demonstrate that they function in the retention of sterols in the PM. Our results have revealed that flippases actually play an essential role in retaining ergosterol in the PM, but the identification of an additional factor, Sfk1, which is totally different from flippases, was essential. Our work indicates that unbiased genetic screening is a powerful approach to understanding cellular mechanisms that are regulated by a different set of proteins. Because Sfk1 is conserved as TMEM150A in mammalian cells [40], cholesterol might be retained in the PM via a similar mechanism.

## Materials and Methods

### Media and chemicals

General chemicals were purchased from Wako Pure Chemicals Industry (Osaka, Japan) unless otherwise stated. Papuamide B was from the collection of R. Andersen. Duramycin was purchased from Sigma-Aldrich (St. Louis, MO, U.S.A). Yeast strains were grown in YPDA-rich medium (1% yeast extract [Difco Laboratories, Detroit, MI, U.S.A.], 2% Bacto-peptone [Difco], 2% glucose, and 0.01% adenine). Strains carrying plasmids were grown in SD synthetic medium (0.67% yeast nitrogen base without amino acids [Difco] and 2% glucose) that contained the required nutritional supplements [111]. The SDA medium was SD medium that contained 0.5% casamino acid (Difco). For the induction of the *GAL1* promoter, 3% galactose and 0.2% sucrose were used as carbon sources (YPGA and SGA-Ura media).

### Yeast strain manipulations and plasmid construction

The yeast strains and plasmids used in this study are listed in S1 and S2 Table, respectively. Standard genetic manipulations of yeast strains were performed according to previously described methods [112]. The polymerase chain reaction (PCR)-based procedure was used to construct yeast strains carrying a complete gene deletion or a gene fusion with either GFP, mCherry, or mRFP1 [113–115]. The amplified DNA fragments were introduced into appropriate strains, and transformants were selected on appropriate plate media. Yeast transformations were performed using the lithium acetate method [116, 117]. All constructs that were produced by the PCR-based procedure were verified by colony PCR to confirm that the replacement or insertion occurred at the expected locus.

When the cell growth phenotype was examined by spot assay, cells were cultured in appropriate medium overnight, adjusted to OD_600_ = 0.64, and then 10-fold serial dilutions were spotted onto the indicated plates.

Strains carrying *3xGFP* or *3xmCherry* at genomic loci were constructed as follows. pBluescript SK+ (pBSK)*-3xGFP-Candida albicans URA3* (*CaURA3*) was constructed by subcloning *3xGFP* from pBSK-*SJL2-3xGFP* [118, 119] (a gift from Drubin, D. G.), *ADH1* terminator, and *CaURA3* into pBSK. Then, a DNA fragment of *SFK1*, *VPH1*, or *DNF3*, which encodes the C-terminal region, was inserted upstream of *3xGFP* in pBSK*-3xGFP-CaURA3*. The resulting plasmids were linearized by cutting at a unique restriction enzyme site in the target gene, followed by transformation into yeast strains. Stable URA^+^ transformants were selected and screened for proper targeting by colony PCR. pBSK-*SFK1-3xmCherry-CaURA3* was constructed by replacing *3xGFP* with *3xmCherry*. After stable *SFK1-3xmCherry::CaURA3* transformants were obtained, *CaURA3* was replaced with the *KanMX6* cassette by marker fragment transformation.

GFP-evt-2PH, GFP-ER (*GFPenvy-SCS2^220-244^*), and *upc2-1* were cloned into pRS306-based vectors and expressed at the *URA3* locus as follows. pRS306-*P_TPI1_*-GFP-evt-2PH-*T_ADH1_* was constructed by replacing mCherry of pRS306-*P_TPI1_*-mCherry-evt-2PH-*T_ADH1_* [120] with GFP. The GFP-ER fragment was generated by PCR with the 3’ primer containing the *SCS2^220-244^* coding region using pColdI-GFPenvy-D4H as a template and inserted into pRS306-*P_TPI1_-T_ADH1_*. The *upc2-1* (G888D) mutant fragment (−800 to +380 bp of the *UPC2* gene) was generated by the standard two-step PCR mutagenesis technique and inserted into pRS306. These plasmids were linearized by cutting at a unique restriction enzyme site in *URA3* and inserted into the *URA3* locus.

To express *OSH* genes on a multicopy plasmid, DNA sequences encoding *OSH* genes were amplified by PCR and subcloned into either YEplac195, YEplac195-*KanMX6*, or YEplac181 plasmids. Sterol binding-deficient *KES1* mutants [72] were generated by the standard two-step PCR mutagenesis technique and subcloned into YEplac195. To express the *SFK1-mCherry* fusion gene on a multicopy plasmid, the *SFK1-mCherry* fragment was generated by overlap extension PCR and subcloned into YEplac181.

To express GFP-D4H and GFPenvy-D4H in *Escherichia coli*, the D4H (D4^D434S^) mutant fragment was generated by the standard two-step PCR mutagenesis technique using pColdI-mCherry-D4 as a template [121]. The GFP and D4H fragments were inserted into pColdI (Takara bio, Shiga, Japan) to construct pColdI-GFP-D4H. To construct pColdI-GFPenvy-D4H, GFPenvy DNA [79] was newly synthesized with codon optimization for *S. cerevisiae* (GeneArt^TM^ Strings, Thermo Scientific, Carlsbad, CA, U.S.A.) and amplified by PCR. pColdI-GFPenvy-D4H was constructed by replacing GFP in pColdI-GFP-D4H with this GFPenvy fragment. To express GFP-D4H and GFPenvy-D4H in yeast, the corresponding DNA fragments were inserted into pRS316-*P_TPI1_-T_ADH1_*.

Schemes detailing the construction of plasmids are available on request.

### Isolation of temperature-sensitive mutations of *SFK1*

The ts *sfk1-2* strain was constructed by PCR-based random mutagenesis as follows. The approximately 1.2 kbp *SFK1* DNA fragment, which corresponds to the region between the 40-bp upstream and 197-bp downstream sequences of the *SFK1* gene, was PCR-amplified under a mutagenic condition [122] using the genomic DNA of the wild-type (YKT38) as a template. On the other hand, the *Kluyveromyces lactis LEU2* (*KlLEU2*) cassette DNA fragment was PCR-amplified under standard conditions using pUG73 (Euroscarf) as a template. In these PCRs, the primers contained additional sequences, so the *SFK1* and *KlLEU2* fragments had overlapping sequences at their 3’ and 5’ regions, respectively. Then, these fragments were used for overlap extension PCR with 5’ (*SFK1*) and 3’ (*KlLEU2*) primers to generate the *SFK1-KILEU2* fragment. This fragment was introduced into YKT2386 (*MAT*a *crf1*Δ*::HphMX4 lem3*Δ*::TRP1 SFK1-GFP::KanMX6*), and the transformants were selected at 30°C for LEU^+^ first and then for G418-sensitive phenotypes. Of these transformants, 265 clones were screened for those that showed growth defects at 37°C. Eight clones were isolated and backcrossed with YKT2332 (*MAT*α *crf1*Δ*::HphMX4 lem3*Δ*::TRP1*) three times. The *crf1*Δ*::HphMX4 lem3*Δ*::TRP1 sfk1-2::KlLEU2*, which which exhibited the tightest ts phenotype, was chosen for further analyses. Sequences of PCR primers used are available on request.

### Isolation of mutants synthetically lethal with the *lem3*Δ *sfk1*Δ mutations

Mutants synthetically lethal with *lem3*Δ *sfk1*Δ were isolated according to the procedures described previously [123]. From 1 × 10^4^ mutagenized cells screened, three single recessive mutations were identified by genetic analyses, and the corresponding wild-type genes were cloned. These genes encode *CRF1*, *DNF3*, and *ANY1*/*CFS1* [124, 125]. Null mutations of these genes were confirmed to be synthetically sick or lethal with *lem3*Δ *sfk1*Δ

### Isolation of multicopy suppressors of the *crf1*Δ *lem3*Δ *sfk1-2* mutant

The *crf1*Δ *lem3*Δ *sfk1-2* mutant (YKT2340) was transformed with a yeast genomic DNA library constructed in the multicopy plasmid YEp24 [126]. Transformants were selected on SDA-Ura plates. The plates were incubated at 25°C for 2 d and then shifted to 37°C, followed by incubation for 3 d. Approximately 1 × 10^6^ transformants were screened, and 186 clones were isolated. To exclude clones that carried *LEM3* or *SFK1*, the sensitivity of the clones to duramycin and cycloheximide was examined [39]. Plasmids were recovered from yeast and reintroduced into the original mutant to confirm the suppression of growth defects. As a result, ten different genomic regions were found to be responsible for suppression by DNA sequencing. The clones that contained a gene relevant to phospholipid asymmetry or lipid metabolism were further analyzed, and *KES1*, *CHO1*, and *CFS1* were identified as suppressors.

### Microscopic observations

For observation of proteins fused to a fluorescent protein in living cells, cells were grown under the indicated conditions to mid-log phase (OD_600_ of 0.8–1.2), collected, mounted on a microslide glass, and immediately observed. Cells were observed under a Nikon ECRIPS E800 microscope (Nikon Instech, Tokyo, Japan) as described previously [14].

Staining of PE exposed to the extracellular leaflet of the PM was performed using the Bio-Ro as described previously [39]. Immunofluorescence staining of Pma1 was performed as described previously [127]. For staining with filipin, cells were grown in YPDA to mid-log phase and fixed with 3.8% formaldehyde for 10 min at room temperature. The fixed cells were washed twice with phosphate-buffered saline (PBS) and resuspended in PBS containing 2.5 mg/mL filipin complex (Sigma-Aldrich). After incubation at room temperature for 15 min in the dark, cells were washed with PBS once and observed with a UV filter set. For TF-Chol labeling, the cells harboring the *upc2-1* mutation were precultured overnight in YPDA and diluted into YPDA containing 0.5% Tween-80, 0.5% ethanol, and 10 µg/mL TF-Chol (Avanti polar lipids, Inc. Alabaster, AL, U.S.A.). The cells were incubated at 30°C for 3 h and then shifted to 37°C, followed by 6 h of incubation. Cells were collected, washed twice with fresh SD medium, resuspended in SD medium, and observed with a GFP filter set. Nile red staining of LDs was performed as described previously with minor modifications [128]. Five OD_600_ units of cell culture were collected and resuspended in 100 µL PBS containing 50 µg/mL Nile red (Sigma-Aldrich). After brief mixing, the cell suspension was incubated for 15 min at room temperature in the dark. Cells were collected, washed five times with PBS, resuspended in PBS, and observed with a G-2A filter set.

Endocytosis was examined by internalization of FM4-64 as described previously with minor modifications [123]. Cells were incubated in YPDA at 30°C for 3 h and then shifted to 37°C, followed by 6 h of incubation. Four OD_600_ units of the cells were labeled with 32 µM FM4-64 (Invitrogen, Madison, WI, U.S.A.) in YPDA on ice for 30 min and then washed once with ice-cold YPDA. Internalization of FM4-64 was initiated by the addition of prewarmed YPDA, and the cells were incubated at 37°C for 30 min, followed by microscopic observation.

### Image analysis

Microscopy image analysis was performed with ImageJ. When the Kes1-GFP-positive abnormal structures were examined, the diameter of the Kes1-GFP signal was measured, and cells with structures larger than approximately 800 nm were classified as having Kes1 abnormal structures.

The PM fluorescence intensity (filipin, Bio-Ro, and Sfk1-mCherry) was analyzed using programmed Macros in ImageJ as follows. (1) The background was subtracted, (2) a cell was selected and its mean fluorescence intensity was quantified (F_whole cell_), (3) the cell periphery (0.2 µm width, 2 pixels) was selected as the PM and its mean fluorescence intensity was quantified (F_pm_), and (4) the signal ratio of the mean F_pm_/F_whole cell_ was calculated.

To analyze the intensity profile of GFPenvy-D4H, the cell periphery was traced with a 2 pixel-wide freehand line tool along the PM of the bud and mother cell. Then, the fluorescence intensity was measured and plotted. The GFPenvy-D4H localization pattern was categorized into three patterns as follows. The PM of budded cells was divided into three regions, the bud PM (bud), the mother cell PM proximal to the bud neck (proximal to bud), and the mother cell PM distal to the bud neck (distal to bud), and the mean fluorescence intensity of each region was calculated as F_bud_, F_proximal to bud_, and F_distal to bud_, respectively. Then, the cells were categorized as “polarized” (F_proximal to bud_ is smaller than 20% of F_bud_), “partially polarized” (F_proximal to bud_ is larger than, but F_distal to bud_ is smaller than, 20% of F_bud_), or “not polarized” (F_distal to bud_ is larger than 20% of F_bud_).

### Rhodamine uptake assay

The rhodamine uptake assay was performed essentially as described previously [39].

### Sucrose density gradient fractionation

Sucrose density gradient fractionation was performed as described previously [13, 87] with minor modifications. Cells were grown at 30°C to mid-log phase in 200 mL YPDA medium and collected. Cells were converted to spheroplasts with zymolyase (Nacalai Tesque, Kyoto, Japan) and broken using a multi-bead shocker (Yasui-Kikai, Osaka, Japan) in break buffer (1.2 M sorbitol, 20 mM HEPES at pH 7.5, 1 mM EDTA, and protease inhibitor cocktail [Nacalai Tesque]). After a 1,000 g spin for 10 min, the resulting supernatant was additionally spun at 13,000 g for 20 min to generate pellet. The step gradient of sucrose was prepared with the following concentrations: 1 mL 60%, 2.5 mL 50%, 2.5 mL 47%, 2 mL 44%, 1 mL 40%, 1 mL 37%, 1 mL 32%, and 0.5 mL 27% (wt/wt) sucrose in a gradient buffer (10 mM HEPES–KOH at pH 7.2 and 1 mM EDTA). The pellet was resuspended in 0.5 mL of the gradient buffer and loaded on the top of the gradient and then centrifuged at 200,000 g in the P40ST rotor (Hitachi, Tokyo, Japan) for 16 h at 4°C. Fractions (0.9 mL) were manually collected from the top of the samples. Pdr5-GFP, Pma1, and Kex2 were detected in each fraction by Western blotting with anti-GFP (Nacalai Tesque), anti-Pma1 (a gift from Serrano R.), and anti-Kex2 antibodies (a gift from Nothwehr S.), respectively.

### Lipid analysis

Cells were grown at 30°C to mid-log phase in 250 mL YPDA medium and collected. Total lipids were extracted by the Bligh and Dyer method [129]. Phospholipid amounts were determined by phosphorus assay [130]. For the phospholipid analysis, samples containing 200 nmol phosphates were subjected to thin-layer chromatography (TLC) plates (Merck, Darmstadt, Germany), and phospholipids were detected as described previously [39]. To detect free and esterified ergosterol, lipid extracts containing 20 nmol phosphates were subjected to high-performance TLC (Merck) separation with hexane/diethyl ether/formic acid (40:10:2, v:v:v). Ergosterols were stained with a mixture of ferric chloride/sulfuric acid/acetic acid by heating [131], and the spots were scanned by an image analyzer. The ergosterol content was determined by TLC-densitometric analysis using ImageJ.

### Liposome sedimentation assay

Recombinant GFP-D4H and GFPenvy-D4H proteins were prepared from *Escherichia coli* as described previously [121]. The protein concentrations were determined by BCA assay.

#### Multilamellar liposomes were prepared by combining

1,2-Dioleoyl-sn-glycero-3-phosphocholine (DOPC, NOF Corporation, Tokyo, Japan) with cholesterol or ergosterol from chloroform stocks. The lipid mixture was evaporated under a stream of nitrogen gas. Then, liposome buffer (0.1 M sucrose, 20 mM HEPES at pH 7.5, 100 mM KCl, and 1 mM EDTA) was added to the dry lipids, and the suspension was vortexed to produce liposomes. D4H binding to liposomes was analyzed as described previously [132] with minor modifications. Recombinant GFP- or GFPenvy-D4H protein (200 nmol) was incubated with liposomes (final total lipid concentration is 100 μM) in HEPES-buffered saline (pH 7.5) for 30 min at room temperature. Then, the mixtures were centrifuged at 21,600 g for 10 min at 25°C. The pellets were washed with HEPES-buffered saline twice. The pellets were subjected to SDS-PAGE followed by Coomassie Brilliant Blue staining. For the quantification of the protein, the stained gel was scanned and analyzed by ImageJ.

### Statistical analysis

To compare the means of multiple groups, statistical analyses were performed using one-way ANOVA followed by Tukey-Kramer multiple comparisons. A p-value <0.05 was regarded as significant. The dataset containing the numerical data and statistical analysis used in this study is listed in S3 Table.

## Supporting information

S1 Table. Saccharomyces cerevisiae strains used in this study.

S2 Table. Plasmids used in this study.

S1 Raw Images. From Figs. 5C, 5D and 7A and S10 Fig.

S3 Table. Dataset containing the numerical data and statistical analysis for this study

## Acknowledgments

We thank Shan Gao for her contribution to the initial stage of this work. We also would like to thank David G Drubin (University of California, Berkeley) and Toshihide Kobayashi (University of Strasbourg) for plasmids, Ramon Serrano (Polytechnic University of Valencia) for the anti-Pma1 antibody, and Steven F Nothwehr (University of Missouri) for the anti-Kex2 antibody. We thank Masato Umeda (Kyoto University) for providing Bio-Ro and Tomohiko Taguchi (Tohoku University) for his helpful comments on evt-2PH. This work was supported by the Japanese Society for the Promotion of Science (JSPS) KAKENHI grants JP18K06104 (T.K.), JP18K14645 (T.M.), and JP19K06536 (K.T.). This work was partly supported by the Photo-excitonix Project at Hokkaido University.

## Supporting Information

**S1 Fig. Growth curves of wild-type and *crf1*Δ *lem3*Δ *skf1-2* cells.**

Cells were precultured in YPDA medium to the mid-log phase at 30°C. Then, the cells were reinoculated in YPDA at time 0 and cultured at 30 or 37°C. Optical density was measured every 1.5 h. Values represent the mean ± S.D. from three independent experiments.

**S2 Fig. Phospholipid distributions in the double mutants of *lem3*Δ, *crf1*Δ, *dnf3*Δ, and *sfk1-2* mutations.**

(A) The *crf1*Δ and *dnf3*Δ mutations increased the sensitivity to PapB and duramycin in the *lem3*Δ mutant. Tenfold serial dilutions of the indicated cell cultures were spotted onto a YPDA plate containing PapB or duramycin, followed by incubation at 30°C for 2 d. (B) GFP -Lact-C2 was normally localized to the PM in the *lem3*Δ *sfk1-2*, *crf1*Δ *sfk1-2*, and *crf1*Δ *lem3*Δ double mutants. Cells were cultured as in Fig 2C. (C) GFP-evt-2PH was normally localized to the PM in the *lem3*Δ *sfk1-2*, *crf1*Δ *sfk1-2*, and *crf1*Δ *lem3*Δ double mutants. Cells were cultured as in Fig 2C. Bars, 5 µm.

**S3 Fig. Analysis of phospholipid composition.**

PC, PI, PS, and PE are quantified as a mol percentage of total phospholipids. The data represent the mean ± S.D. derived from the analysis of four to five independent samples. “n.s,” indicates no significant difference between all combinations as determined by the Tukey–Kramer test.

**S4 Fig. Nutrient transporters fail to localize to the PM in the triple but not in the double mutants.**

(A) Can1-GFP (upper panel) and Hip1-GFP (lower panel) were localized to the PM in the *lem3*Δ *sfk1-2*, *crf1*Δ *sfk1-2*, and *crf1*Δ *lem3*Δ double mutants. Cells were cultured as in Fig 2C. (B) Nutrient transporters failed to localize to the PM in the *crf1*Δ *lem3*Δ *sfk1-2* triple mutant. Amino acid transporters (Alp1-, Lyp1-, Tat1-, and Ptr2-GFP) and glucose transporters (Hxt2-, 3-, and 4-GFP) were examined. Cells were cultured as in Fig 2C. (C) Can1-GFP and Hip1-GFP were mislocalized to the vacuole by endocytosis in the *crf1*Δ *lem3*Δ *sfk1-2* triple mutant. Cells were cultured in YPDA at 37°C for 5.5 h, followed by additional incubation for 30 min in the presence (LAT-A) or absence (DMSO) of 100 µM LAT-A. *Right panel*: the percentage of cells with Can1-GFP or Hip1-GFP at the PM was determined and is expressed as the mean ± S.D. of three independent experiments (n>168 cells in total for each strain). Asterisks indicate a significant difference, as determined by the Tukey–Kramer test (p < 0.05). Bars, 5 µm.

**S5 Fig. The defects in the *crf1*Δ *lem3*Δ *sfk1-2* mutant may be independent of PI(4)P.**

(A) The distribution of the Osh2-PH-GFP PI(4)P biosensor was not altered in the *crf1*Δ *lem3*Δ *sfk1-2* triple mutant. Cells were cultured as in Fig 2C. Bar, 5 µm. *Right panel*: The localization of Osh2-PH-GFP was categorized into four patterns. The percentages of cells with these patterns were determined and expressed as the mean ± S.D. of three independent experiments (n > 142 cells in total for each strain). “n.s,” indicates no significant difference between all combinations as determined by the Tukey–Kramer test. (B) The C-terminal truncation of *SFK1* did not affect the growth of the *crf1*Δ *lem3*Δ double mutant. The *SFK1ΔC-GFP* mutant in which the C-terminal cytoplasmic region was deleted [39] was combined with *crf1*Δ *lem3*Δ mutations. Tenfold serial dilutions were spotted onto a YPDA plate, followed by incubation for 1.5 d at 30 or 37°C.

**S6 Fig. Normal localizations of Kes1-GFP and GFP-ER in the *lem3*Δ *sfk1-2*, *crf1*Δ *sfk1-2*, and *crf1*Δ *lem3*Δ double mutants.**

The localizations of Kes1-GFP (A) and GFP-ER (B) in the double mutants are shown. Cells were cultured as in Fig 2C. Bars, 5 µm.

**S7 Fig. Filipin staining in the *lem3***Δ ***sfk1***Δ**, *crf1***Δ ***sfk1***Δ**, and *crf1***Δ ***lem3***Δ **double mutants.**

Cells were cultured and stained with filipin as in Fig 5A. Bar, 5 µm.

**S8 Fig. Ergosterol-dependent PM localization of GFPenvy-D4H.**

(A) Fluconazole treatment inhibits the GFPenvy-D4H distribution to the PM. Wild-type cells harboring pRS316-GFPenvy-D4H were grown in SDA-Ura medium to the mid-log phase at 30°C and then treated with 100 µM fluconazole or mock treated, followed by incubation for 6 h at 30°C. *Right panel*: the percentage of cells with GFPenvy-D4H at the PM was determined and is expressed as the mean ± S.D. of three independent experiments (n > 257 cells in total for each condition). An asterisk indicates a significant difference, as determined by the Tukey–Kramer test (p < 0.05). (B) The distribution of GFPenvy-D4H in mutants of genes encoding the enzymes in the late steps of ergosterol biosynthesis (*ERG2-6*). Cells were cultured in SDA-Ura medium at 30°C to the mid-log phase. (C) GFPenvy-D4H was localized to the PM in the *lem3*Δ *sfk1-2*, *crf1*Δ *sfk1-2*, and *crf1Δ lem3*Δ double mutants. Cells were cultured as in Fig 2C, except that SDA-Ura medium was used. (D) The PM localization of GFPenvy-D4H was partially recovered by *KES1* overexpression in the *crf1*Δ *lem3*Δ *sfk1-2* triple mutant. Cells harboring pRS316-GFPenvy-D4H and either YEplac181-*KES1* or YEplac181 were cultured as in Fig 2C except that SD-Leu-Ura medium was used. Arrows indicate the PM localization of GFPenvy-D4H. *Right panel*: the percentage of cells with GFPenvy-D4H at the PM was determined and is expressed as the mean ± S.D. of three independent experiments (n > 121 in total for each strain). An asterisk indicates a significant difference, as determined by the Tukey–Kramer test (p < 0.05). Bars, 5 µm.

**S9 Fig. TF-Chol is retained in the PM of the *lem3Δ sfk1-2*, *crf1Δ sfk1-2*, and *crf1*Δ *lem3*Δ double mutants.**

Cells were cultured and labeled with TF-Chol as described in the “Materials and Methods”. Bar, 5 µm.

**S10 Fig. Identification of esterified ergosterol.**

TLC analysis of total sterols was performed as in Fig 7A. To detect esterified ergosterol, total lipids were extracted from cells in the stationary phase.

**S11 Fig. Lipid droplets are increased in the *crf1*Δ *lem3*Δ *sfk1-2* triple mutant.**

(A) Nile red staining in the *lem3*Δ *sfk1*Δ, *crf1*Δ *sfk1*, and *crf1*Δ *lem3*Δ double mutants. Cells were cultured in YPDA medium to the mid-log phase at 30°C, followed by Nile red staining. Nile red staining was performed as described in the “Materials and Methods”. (B) Accumulation of Tgl1-GFP (left) and Faa4-GFP (right) puncta in the *crf1*Δ *lem3*Δ *sfk1-2* triple mutant. Cells were cultured as in Fig 2C. All images were acquired and processed under the same conditions for comparison of fluorescence intensity. Bars, 5 µm.

**S12 Fig. The growth defect in the *crf1*Δ *lem3*Δ *sfk1-2* triple mutant is independent of triacylglycerol.**

(A) The growth defect of the *crf1*Δ *lem3*Δ *sfk1-2* triple mutant is not suppressed by either the *dga1*Δ or *lro1*Δ mutation. Tenfold serial dilutions were spotted onto a YPDA plate, followed by incubation for 1.5 d at 30 or 37°C. (B) Synthetic growth defects in the *are2*Δ *upc2-1* mutant. Diploid cells with the indicated genotype were sporulated, dissected, and grown at 30°C for 4 d. Tetrad genotypes were determined as in Fig 1B, and the identities of the double mutant segregants are shown in parentheses (red circles).

**S13 Fig. Classification of Sfk1-mCherry-expressing cells with low or high expression patterns.**

(A) Fluorescence intensity profile of a cell showing the “partially polarized” pattern of GFPenvy-D4H. Fluorescence signals were quantified along the dotted line from the mother cell to the bud. The brightness of Sfk1-3xmCherry was adjusted as in Fig 9B. Bar, 3 µm. (B) Classification of Sfk1-mCherry-expressing cells with low or high expression patterns. The cells in Fig 9E were examined. Low or high expression of Sfk1-mCherry was determined for each cell on the basis of our threshold value, which was set at 300% of the fluorescence intensity of endogenously expressed Sfk1-3xmCherry. Fluorescence intensity at the PM was quantitated as described in the “Materials and Methods”. The ratio of the fluorescence at the PM (F_pm_)/that of whole cell (F_whole cell_) was determined and expressed with a boxplot (whiskers: maximum and minimum values; box: first quartile, median, and third quartile; circle: average). *Bars*: Endo, endogenous expression of Sfk1-3xmCherry; Low, multicopy plasmid of *SFK1-mCherry* but low expression of Sfk1-mCherry; High, multicopy plasmid of *SFK1-mCherry* and high expression of Sfk1-mCherry. The numbers of cells analyzed were 13, 34, and 22 for Endo, Low, and High, respectively. An asterisk indicates a significant difference, as determined by the Tukey–Kramer test (p < 0.05).

**S1 Raw Images. Raw images underlying figures.**

From Figs. 5C, 5D and 7A and S10 Fig.

**S1 Table. *Saccharomyces cerevisiae* strains used in this study.**

**S2 Table. Plasmids used in this study.**

**S3 Table. Excel spreadsheet containing the numerical data and statistical analysis for Figs 2E, 2F, 3A, 3C, 3D, 4B, 4C, 4E, 5A, 5B, 5C, 5D, 5E, 5F, 6B, 7A, 7B, 7C, 8B, 9C, 9D, 9F, and 9G, S1, S3, S4C, S5A, S8A, S8D, S13A and S13B Figs.**

## Notes

### Competing Interest Statement

The authors have declared no competing interest.

