## Supplementary material for "Phospholipid flippases and Sfk1 are essential for the retention of ergosterol in the plasma membrane": S1-13 Fig. Supporting Figs.

S1 Fig

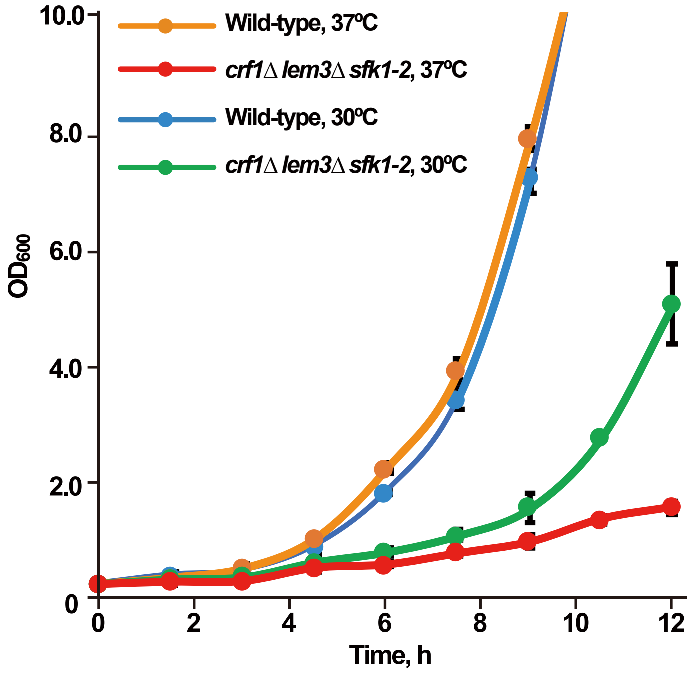

Wild-type, 37ºC
crf1Δ lem3Δ sfk1-2, 37ºC
Wild-type, 30ºC
crf1Δ lem3Δ sfk1-2, 30ºC
10.0
8.0
6.0
4.0
2.0
0
OD600
0
2
4
6
8
10
12
Time, h

S2 Fig

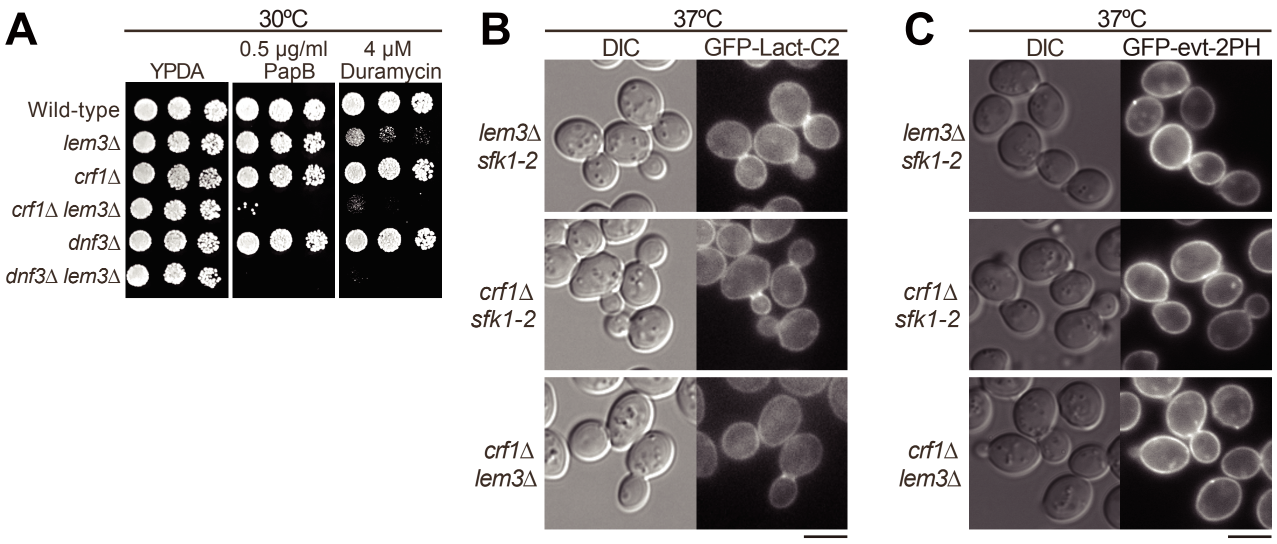
A
30ºC
YPDA
0.5 µg/ml PapB
4 µM Duramycin
Wild-type
lem3Δ
crf1Δ
crf1Δ lem3Δ
dnf3Δ
dnf3Δ lem3Δ
B
37ºC
DIC
GFP-Lact-C2
lem3Δ sfk1-2
crf1Δ sfk1-2
crf1Δ lem3Δ
C
37ºC
DIC
GFP-evt-2PH
lem3Δ sfk1-2
crf1Δ sfk1-2
crf1Δ lem3Δ

S3 Fig

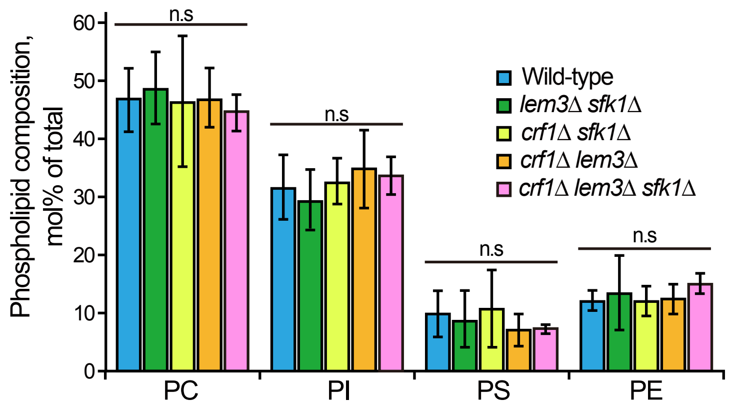

Phospholipid composition, mol% of total
60
50
40
30
20
10
0
n.s
n.s
n.s
n.s
Wild-type
lem3Δ sfk1Δ
crf1Δ sfk1Δ
crf1Δ lem3Δ
crf1Δ lem3Δ sfk1Δ
PC
PI
PS
PE

S4 Fig

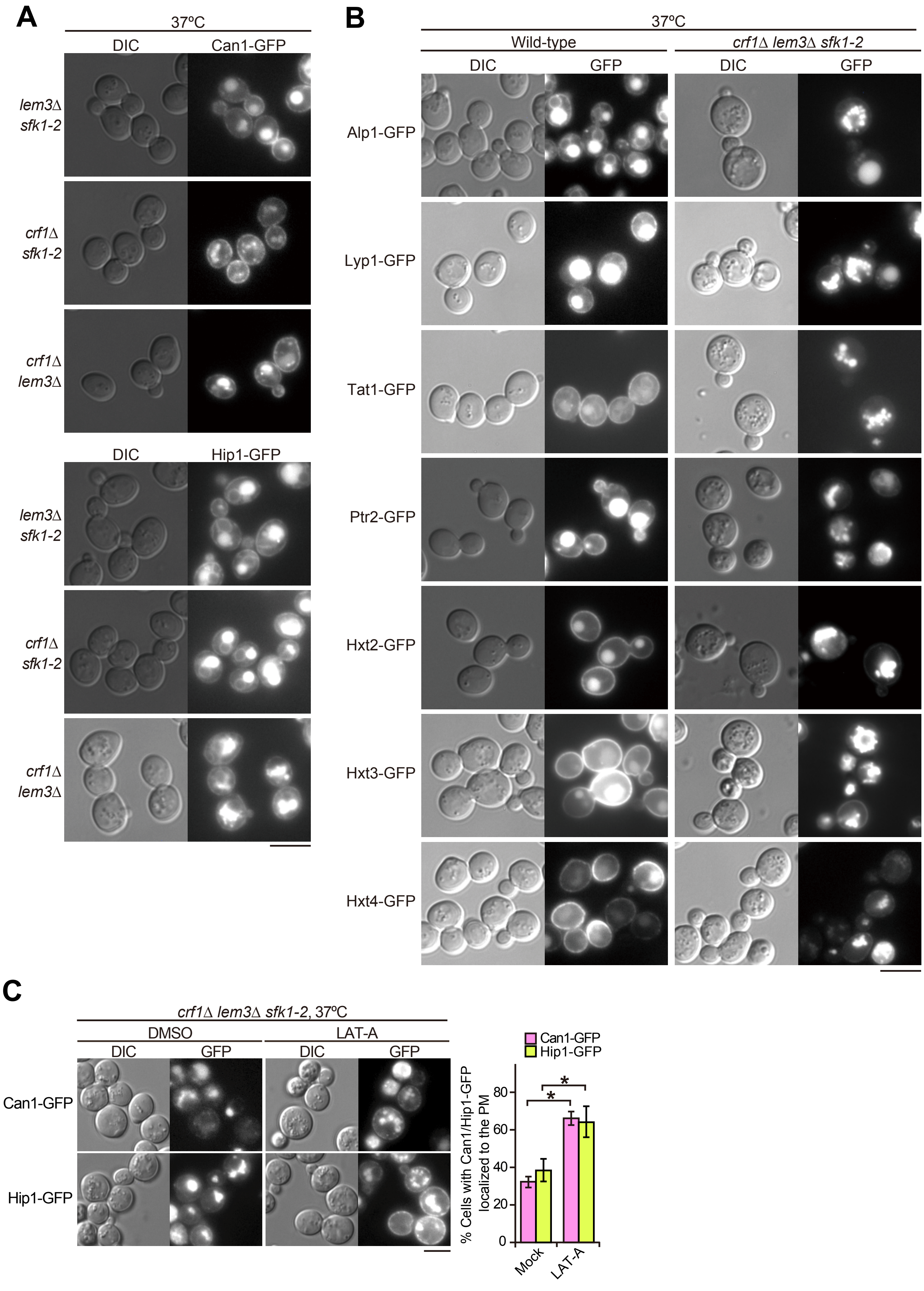
A
37ºC
DIC
Can1-GFP
lem3Δ sfk1-2
crf1Δ sfk1-2
crf1Δ lem3Δ
DIC
Hip1-GFP
lem3Δ sfk1-2
crf1Δ sfk1-2
crf1Δ lem3Δ
B
37ºC
Wild-type
crf1Δ lem3Δ sfk1-2
DIC
GFP
DIC
GFP
Alp1-GFP
Lyp1-GFP
Tat1-GFP
Ptr2-GFP
Hxt2-GFP
Hxt3-GFP
Hxt4-GFP
C
crf1Δ lem3Δ sfk1-2, 37ºC
DMSO
LAT-A
DIC
GFP
DIC
GFP
Can1-GFP
Hip1-GFP
Can1-GFP
Hip1-GFP
% Cells with Can1/Hip1-GFP localized to the PM
0
20
40
60
80
*
*
Mock
LAT-A

S5 Fig

A

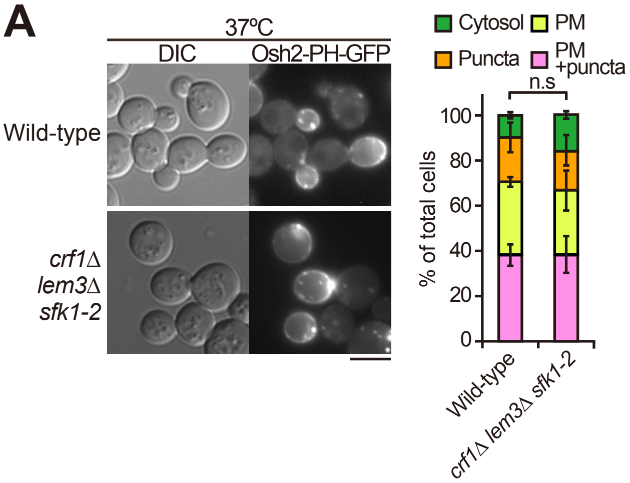
37ºC
DIC
Osh2-PH-GFP
Wild-type
crf1Δ lem3Δ sfk1-2
Cytosol
PM
Puncta
PM +puncta
n.s
% of total cells
100
80
60
40
20
0
Wild-type
crf1Δ lem3Δ sfk1-2

B

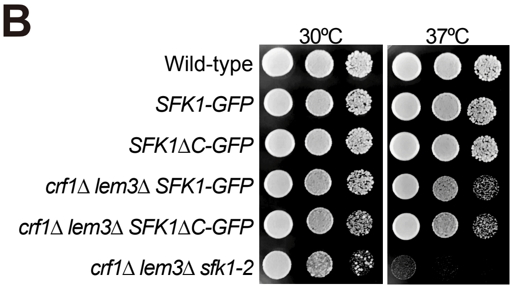
30ºC
37ºC
Wild-type
SFK1-GFP
SFK1ΔC-GFP
crf1Δ lem3Δ SFK1-GFP
crf1Δ lem3Δ SFK1ΔC-GFP
crf1Δ lem3Δ sfk1-2

S6 Fig

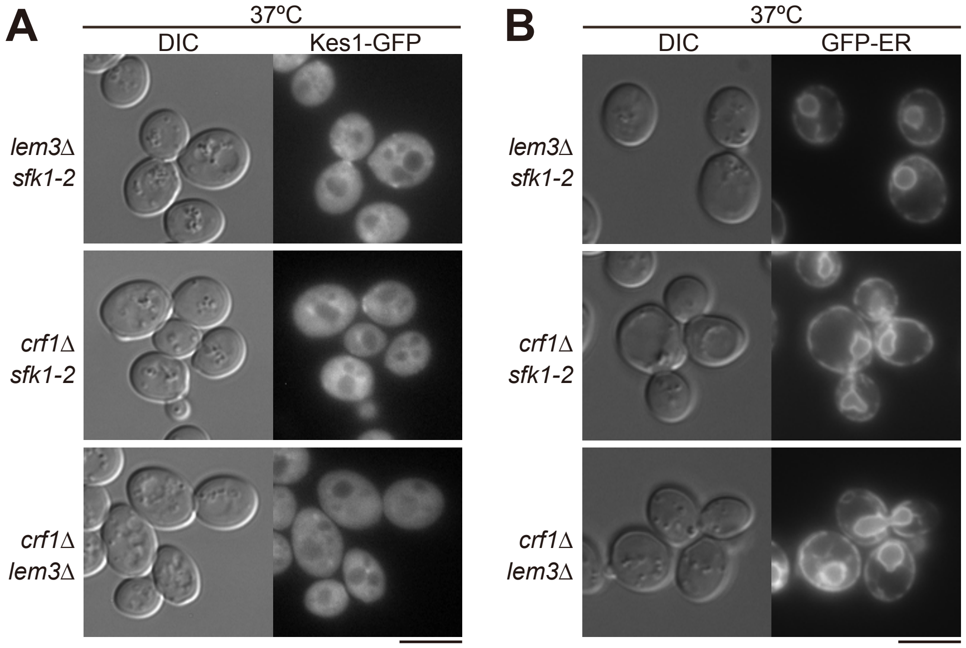
A
37ºC
DIC
Kes1-GFP
lem3Δ sfk1-2
crf1Δ sfk1-2
crf1Δ lem3Δ
B
37ºC
DIC
GFP-ER
lem3Δ sfk1-2
crf1Δ sfk1-2
crf1Δ lem3Δ

### S7 Fig

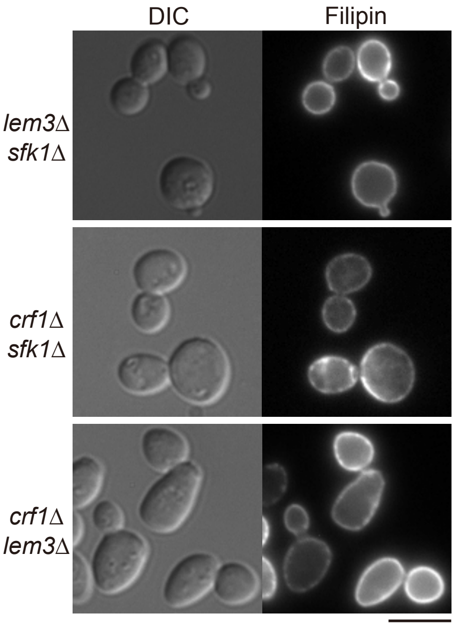

### S8 Fig

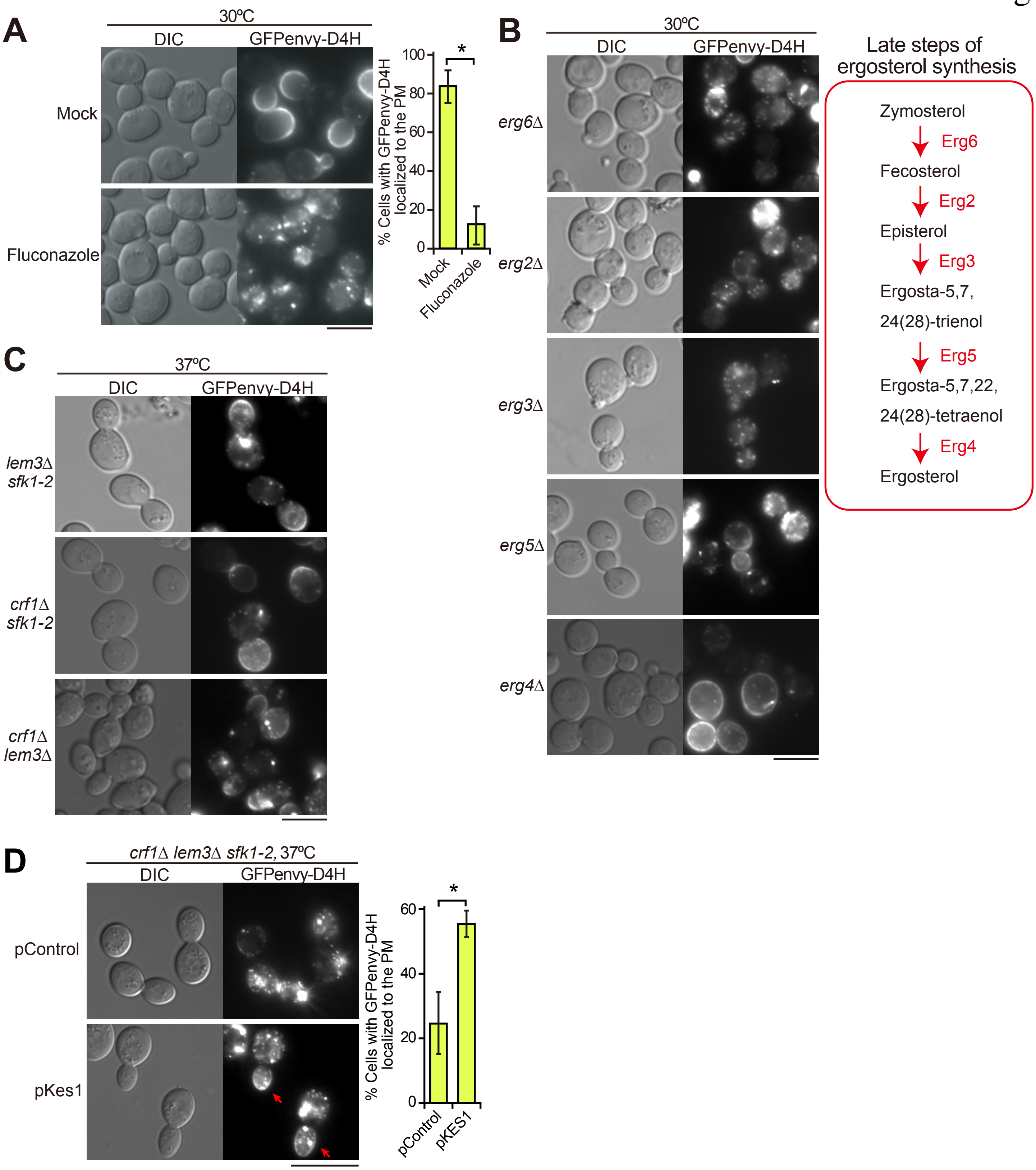

A
30ºC
DIC
GFPenvy-D4H
Mock
Fluconazole
% Cells with GFPenvy-D4H localized to the PM
100
80
60
40
20
0
*
Mock
Fluconazole
B
30ºC
DIC
GFPenvy-D4H
erg6Δ
erg2Δ
erg3Δ
erg5Δ
erg4Δ
Late steps of ergosterol synthesis
Zymosterol
Erg6
Fecosterol
Erg2
Episterol
Erg3
Ergosta-5,7,
24(28)-trienol
Erg5
Ergosta-5,7,22,
24(28)-tetraenol
Erg4
Ergosterol
C
37ºC
DIC
GFPenvy-D4H
lem3Δ sfk1-2
crf1Δ sfk1-2
crf1Δ lem3Δ
D
crf1Δ lem3Δ sfk1-2, 37ºC
DIC
GFPenvy-D4H
pControl
pKes1
% Cells with GFPenvy-D4H localized to the PM
60
40
20
0
*
pControl
pKES1

S9 Fig

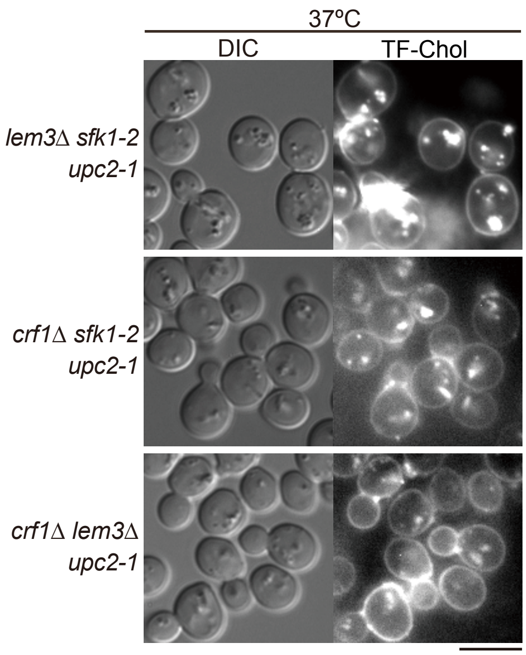
37ºC
DIC
TF-Chol
lem3Δ sfk1-2
upc2-1
crf1Δ sfk1-2
upc2-1
crf1Δ lem3Δ
upc2-1

S10 Fig

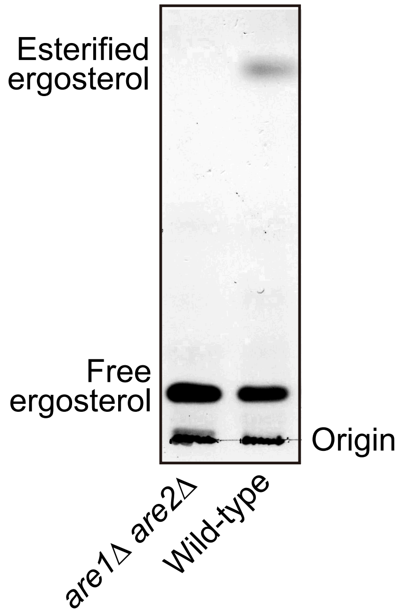
Esterified ergosterol
Free ergosterol
Origin
*are1Δ are2Δ*
Wild-type

S11 Fig

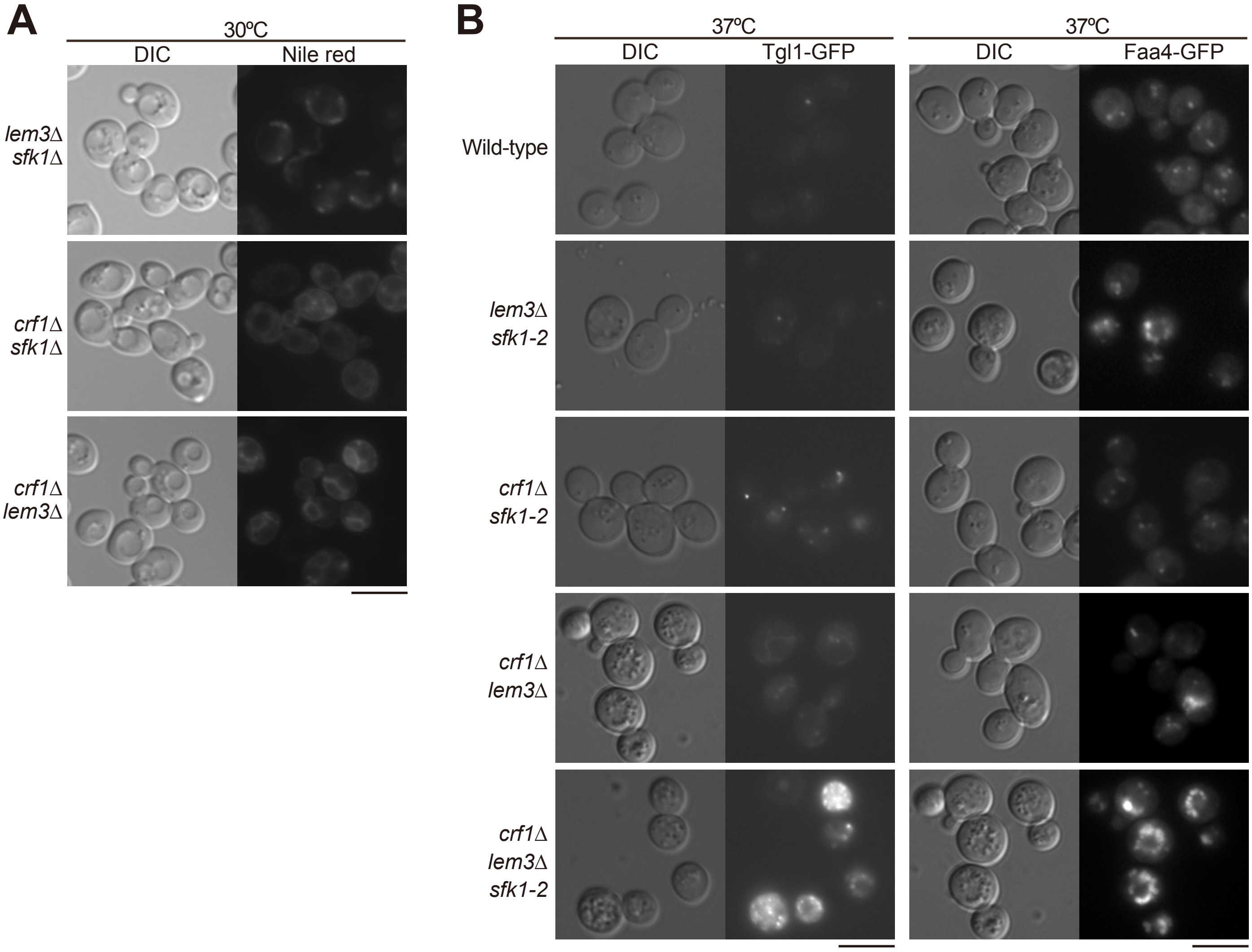
A
30ºC
DIC
Nile red
lem3Δ sfk1Δ
crf1Δ sfk1Δ
crf1Δ lem3Δ
B
37ºC
DIC
Tgl1-GFP
37ºC
DIC
Faa4-GFP
Wild-type
lem3Δ sfk1-2
crf1Δ sfk1-2
crf1Δ lem3Δ
crf1Δ lem3Δ sfk1-2

S12 Fig

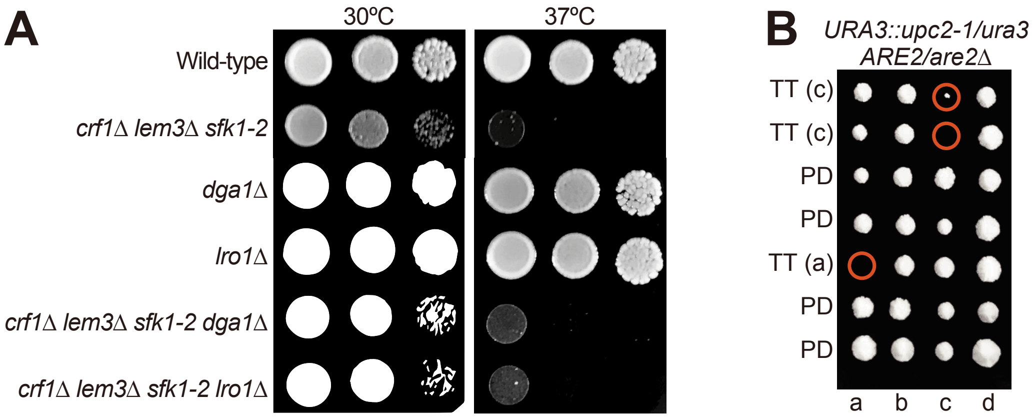
A
30ºC
37ºC
Wild-type
crf1Δ lem3Δ sfk1-2
dga1Δ
lro1Δ
crf1Δ lem3Δ sfk1-2 dga1Δ
crf1Δ lem3Δ sfk1-2 lro1Δ
B
URA3::upc2-1/ura3
ARE2/are2Δ
TT (c)
TT (c)
PD
PD
TT (a)
PD
PD
a
b
c
d

### S13 Fig

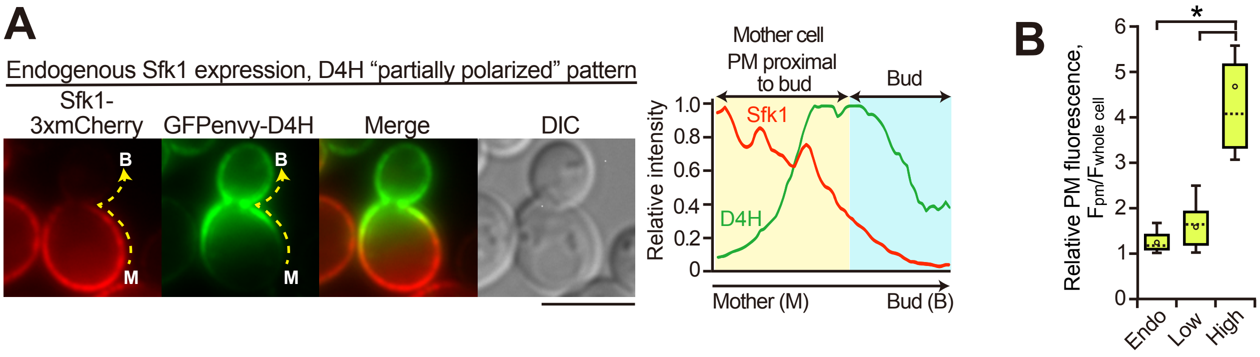

A
Endogenous Sfk1 expression, D4H "partially polarized" pattern
Sfk1-3xmCherry
GFPenvy-D4H
Merge
DIC
B
M
Mother cell
PM proximal to bud
Bud
Relative intensity
Sfk1
D4H
Mother (M)
Bud (B)
B
Relative PM fluorescence, $F_{pm}/F_{whole\ cell}$
*
Endo
Low
High
