## Supplementary material for "Phospholipid flippases and Sfk1 are essential for the retention of ergosterol in the plasma membrane": S1 Table. Saccharomyces cerevisiae strains used in this study.

**S1 Table.** *S. cerevisiae* strains used in this study

| Strain[a,b] | Mating type | Genotype | Reference or source |
|---|---|---|---|
| YKT38 | a | Wild-type (*lys2-801 ura3-52 his3Δ-200 trp1Δ-63 leu2Δ-1*) | [1] |
| YKT715 | a | *lem3Δ::TRP1* | [2] |
| YKT2329 | a | *crf1Δ::HphMX4* | This study |
| YKT2117 | a | *sfk1Δ::KanMX6* | [3] |
| YKT2330 | a | *lem3Δ::TRP1 sfk1Δ::KanMX6* | This study |
| YKT2331 | a | *crf1Δ::HphMX4 sfk1Δ::KanMX6* | This study |
| YKT2332 | α | *crf1Δ::HphMX4 lem3Δ::TRP1* | This study |
| YKT2333 | a | *crf1Δ::HphMX4 lem3Δ::TRP1 sfk1Δ::KanMX6* | This study |
| YKT2334 | α | *dnf3Δ::HIS3MX6 sfk1Δ::KanMX6* | This study |
| YKT783 | a | *CRF1-GFP::KanMX6* | This study |
| YKT1412 | α | *vrp1Δ::LUE2 CRF1-GFP::KanMX6* | This study |
| YKT2335 | a | *DNF3-3xGFP::CaURA3* | This study |
| YKT2336 | a | *vrp1Δ::LUE2 DNF3-3xGFP::CaURA3* | This study |
| YKT2337 | a | *sfk1-2::KlLEU2* | This study |
| YKT2338 | a | *lem3Δ::TRP1 sfk1-2::KlLEU2* | This study |
| YKT2339 | a | *crf1Δ::HphMX4 sfk1-2::KlLEU2* | This study |
| YKT2340 | a | *crf1Δ::HphMX4 lem3Δ::TRP1 sfk1-2::KlLEU2* | This study |
| YKT917 | α | *dnf3Δ::HIS3MX6* | This study |
| YKT2341 | a | *dnf3Δ::HIS3MX6 lem3Δ::TRP1* | This study |
| YKT2235 | a | *SFK1-3xGFP::CaURA3* | This study |
| YKT2342 | a | *sfk1-2-3xGFP::CaURA3::KlLEU2* | This study |
| YKT2343 | a | *URA3::GFP-Lact-C2* | [4] |
| YKT2344 | a | *lem3Δ::TRP1 sfk1-2::KlLEU2 URA3::GFP-Lact-C2* | This study |
| YKT2345 | a | *crf1Δ::HphMX4 sfk1-2::KlLEU2 URA3::GFP-Lact-C2* | This study |
| YKT2346 | a | *crf1Δ::HphMX4 lem3Δ::TRP1 URA3::GFP-Lact-C2* | This study |
| YKT2347 | a | *crf1Δ::HphMX4 lem3Δ::TRP1 sfk1-2::KlLEU2 URA3::GFP-Lact-C2* | This study |
| YKT2348 | a | *URA3::GFP-evt-2PH* | This study |
| YKT2349 | a | *lem3Δ::TRP1 sfk1-2::KlLEU2 URA3::GFP-evt-2PH* | This study |
| YKT2350 | a | *crf1Δ::HphMX4 sfk1-2::KlLEU2 URA3::GFP-evt-2PH* | This study |
| YKT2351 | a | *crf1Δ::HphMX4 lem3Δ::TRP1 URA3::GFP-evt-2PH* | This study |
| YKT2352 | a | *crf1Δ::HphMX4 lem3Δ::TRP1 sfk1-2::KlLEU2 URA3::GFP-evt-2PH* | This study |

*Continued*

**Table S1.** *Continued*

| Strain[a,b] | Mating type | Genotype | Reference or source |
|---|---|---|---|
| YKT2354 | a | *URA3::GFP-SNC1* | [5] |
| YKT2355 | a | *crf1Δ::HphMX4 lem3Δ::TRP1 sfk1-2::KlLEU2 URA3::GFP-SNC1* | This study |
| YKT1187 | a | *PDR5-GFP::KanMX6* | This study |
| YKT2356 | a | *crf1Δ::HphMX4 lem3Δ::TRP1 sfk1-2::KlLEU2 PDR5-GFP::KanMX6* | This study |
| YKT2357 | a | *VPH1-3xGFP::CaURA3* | This study |
| YKT2358 | a | *crf1Δ::HphMX4 lem3Δ::TRP1 sfk1-2::KlLEU2 VPH1-3xGFP::CaURA3* | This study |
| YKT2359 | a | *CAN1-GFP::CaURA3* | This study |
| YKT2360 | a | *lem3Δ::TRP1 sfk1-2::KlLEU2 CAN1-GFP::CaURA3* | This study |
| YKT2361 | a | *crf1Δ::HphMX4 sfk1-2::KlLEU2 CAN1-GFP::CaURA3* | This study |
| YKT2362 | a | *crf1Δ::HphMX4 lem3Δ::TRP1 CAN1-GFP::CaURA3* | This study |
| YKT2363 | a | *crf1Δ::HphMX4 lem3Δ::TRP1 sfk1-2::KlLEU2 CAN1-GFP::CaURA3* | This study |
| YKT2364 | a | *HIP1-GFP::CaURA3* | This study |
| YKT2365 | a | *lem3Δ::TRP1 sfk1-2::KlLEU2 HIP1-GFP::CaURA3* | This study |
| YKT2366 | a | *crf1Δ::HphMX4 sfk1-2::KlLEU2 HIP1-GFP::CaURA3* | This study |
| YKT2367 | a | *crf1Δ::HphMX4 lem3Δ::TRP1 HIP1-GFP::CaURA3* | This study |
| YKT2368 | a | *crf1Δ::HphMX4 lem3Δ::TRP1 sfk1-2::KlLEU2 HIP1-GFP::CaURA3* | This study |
| YKT2369 | a | *ALP1-GFP::CaURA3* | This study |
| YKT2370 | a | *crf1Δ::HphMX4 lem3Δ::TRP1 sfk1-2::KlLEU2 ALP1-GFP::CaURA3* | This study |
| YKT2371 | a | *LYP1-GFP::CaURA3* | This study |
| YKT2372 | a | *crf1Δ::HphMX4 lem3Δ::TRP1 sfk1-2::KlLEU2 LYP1-GFP::CaURA3* | This study |
| YKT2373 | a | *TAT1-GFP::CaURA3* | This study |
| YKT2374 | a | *crf1Δ::HphMX4 lem3Δ::TRP1 sfk1-2::KlLEU2 TAT1-GFP::CaURA3* | This study |
| YKT2375 | a | *PTR2-GFP::CaURA3* | This study |
| YKT2376 | a | *crf1Δ::HphMX4 lem3Δ::TRP1 sfk1-2::KlLEU2 PTR2-GFP::CaURA3* | This study |
| YKT2377 | a | *HXT2-GFP::CaURA3* | This study |
| YKT2378 | a | *crf1Δ::HphMX4 lem3Δ::TRP1 sfk1-2::KlLEU2 HXT2-GFP::CaURA3* | This study |
| YKT2379 | a | *HXT3-GFP::CaURA3* | This study |
| YKT2380 | a | *crf1Δ::HphMX4 lem3Δ::TRP1 sfk1-2::KlLEU2 HXT3-GFP::CaURA3* | This study |
| YKT2381 | a | *HXT4-GFP::CaURA3* | This study |
| YKT2382 | a | *crf1Δ::HphMX4 lem3Δ::TRP1 sfk1-2::KlLEU2 HXT4-GFP::CaURA3* | This study |
| YKT2251 | α | *URA3::OSH2-2xPH-3xGFP* (*OSH2-PH-GFP*) | [6] |

*Continued*

**Table S1.** *Continued*

| Strain[a,b] | Mating type | Genotype | Reference or source |
|---|---|---|---|
| YKT2383 | a | *crf1Δ::HphMX4 lem3Δ::TRP1 sfk1*-2*::KlLEU2 URA3::OSH2-2xPH-3xGFP* | This study |
| YKT2120 | α | *SFK1-GFP::KanMX6* | [3] |
| YKT2385 | a | *SFK1ΔC-GFP::KanMX6* | This study |
| YKT2386 | a | *crf1Δ::HphMX4 lem3Δ::TRP1 SFK1-GFP::KanMX6* | This study |
| YKT2387 | a | *crf1Δ::HphMX4 lem3Δ::TRP1 SFK1ΔC-GFP::KanMX6* | This study |
| YKT2388 | a | *KES1-GFP::CaURA3* | This study |
| YKT2389 | a | *lem3Δ::TRP1 sfk1*-2*::KlLEU2 KES1-GFP::CaURA3* | This study |
| YKT2390 | a | *crf1Δ::HphMX4 sfk1*-2*::KlLEU2 KES1-GFP::CaURA3* | This study |
| YKT2391 | a | *crf1Δ::HphMX4 lem3Δ::TRP1 KES1-GFP::CaURA3* | This study |
| YKT2392 | a | *crf1Δ::HphMX4 lem3Δ::TRP1 sfk1*-2*::KlLEU2 KES1-GFP::CaURA3* | This study |
| YKT2393 | a | *URA3::*$P_{TPI1}$*-GFPenvy-SCS2*$^{220\text{-}244}$ (*GFP-ER*) | This study |
| YKT2394 | a | *lem3Δ::TRP1 sfk1*-2*::KlLEU2 URA3::*$P_{TPI1}$*-GFPenvy-SCS2*$^{220\text{-}244}$ | This study |
| YKT2395 | a | *crf1Δ::HphMX4 sfk1*-2*::KlLEU2 URA3::*$P_{TPI1}$*-GFPenvy-SCS2*$^{220\text{-}244}$ | This study |
| YKT2396 | a | *crf1Δ::HphMX4 lem3Δ::TRP1 URA3::*$P_{TPI1}$*-GFPenvy-SCS2*$^{220\text{-}244}$ | This study |
| YKT2397 | a | *crf1Δ::HphMX4 lem3Δ::TRP1 sfk1*-2*::KlLEU2 URA3::*$P_{TPI1}$*-GFPenvy-SCS2*$^{220\text{-}244}$ | This study |
| YKT2398 | a | *KES1-GFP::CaURA3 SEC63-mRFP1::KanMX6* | This study |
| YKT2399 | a | *crf1Δ::HphMX4 lem3Δ::TRP1 sfk1*-2*::KlLEU2 KES1-GFP::CaURA3 SEC63-mRFP1::KanMX6* | This study |
| YKT2400 | a | *HIS3MX6::*$P_{GAL1}$*-3HA-ERG11* | This study |
| YKT2119 | a | *erg6Δ::KanMX6 TRP1* | [3] |
| YKT2401 | a | *erg2Δ::TRP1* | This study |
| KKT12 | a | *erg3Δ::HphMX4 TRP1* | [7] |
| YKT2402 | a | *erg5Δ::HIS3MX6* | This study |
| KKT252 | a | *erg4Δ::HphMX4 TRP1* | [7] |
| YKT2403 | α | *URA3::upc2-1* | This study |
| YKT2404 | a | *lem3Δ::TRP1 sfk1*-2*::NatMX6 URA3::upc2-1* | This study |
| YKT2405 | a | *crf1Δ::HphMX4 sfk1*-2*::NatMX6 URA3::upc2-1* | This study |
| YKT2406 | a | *crf1Δ::HphMX4 lem3Δ::TRP1 URA3::upc2-1* | This study |
| YKT2407 | a | *crf1Δ::HphMX4 lem3Δ::TRP1 sfk1*-2*::NatMX6 URA3::upc2-1* | This study |
| YKT2408 | a | *TGL1-GFP::CaURA3* | This study |

*Continued*

**Table S1.** *Continued*

| Strain[a,b] | Mating type | Genotype | Reference or source |
|---|---|---|---|
| YKT2409 | a | *lem3Δ::TRP1 sfk1-2::KlLEU2 TGL1-GFP::CaURA3* | This study |
| YKT2410 | a | *crf1Δ::HphMX4 sfk1-2::KlLEU2 TGL1-GFP::CaURA3* | This study |
| YKT2411 | a | *crf1Δ::HphMX4 lem3Δ::TRP1 TGL1-GFP::CaURA3* | This study |
| YKT2412 | a | *crf1Δ::HphMX4 lem3Δ::TRP1 sfk1-2::KlLEU2 TGL1-GFP::CaURA3* | This study |
| YKT2413 | a | *FAA4-GFP::CaURA3* | This study |
| YKT2414 | a | *lem3Δ::TRP1 sfk1-2::KlLEU2 FAA4-GFP::CaURA3* | This study |
| YKT2415 | a | *crf1Δ::HphMX4 sfk1-2::KlLEU2 FAA4-GFP::CaURA3* | This study |
| YKT2416 | a | *crf1Δ::HphMX4 lem3Δ::TRP1 FAA4-GFP::CaURA3* | This study |
| YKT2417 | a | *crf1Δ::HphMX4 lem3Δ::TRP1 sfk1-2::KlLEU2 FAA4-GFP::CaURA3* | This study |
| YKT2418 | a | *FAA4-mCherry::KanMX6 URA3::upc2-1* | This study |
| YKT2419 | a | *crf1Δ::HphMX4 lem3Δ::TRP1 sfk1-2::KlLEU2 FAA4-mCherry::KanMX6 URA3::upc2-1* | This study |
| YKT2420 | a | *are1Δ::KanMX6* | This study |
| YKT2421 | a | *are2Δ::HIS3MX6* | This study |
| YKT2422 | a | *crf1Δ::HphMX4 lem3Δ::TRP1 sfk1-2::KlLEU2 are1Δ::KanMX6* | This study |
| YKT2423 | a | *crf1Δ::HphMX4 lem3Δ::TRP1 sfk1-2::KlLEU2 are2Δ::HIS3MX6* | This study |
| YKT2424 | a | *dga1Δ::HIS3MX6* | This study |
| YKT2425 | a | *lro1Δ::KanMX6* | This study |
| YKT2426 | a | *crf1Δ::HphMX4 lem3Δ::TRP1 sfk1-2::KlLEU2 dga1Δ::HIS3MX6* | This study |
| YKT2427 | a | *crf1Δ::HphMX4 lem3Δ::TRP1 sfk1-2::KlLEU2 lro1Δ::KanMX6* | This study |
| YKT2428 | a | *SFK1-3xmCherry::KanMX6* | This study |

[a] YKT strains are isogenic derivatives of YEF473 [8]. Only relevant genotypes are described.
[b] KKT12 and KKT252 strains are isogenic derivatives of BY4743[9].

2. Saito K, Fujimura-Kamada K, Furuta N, Kato U, Umeda M, Tanaka K. Cdc50p,

a protein required for polarized growth, associates with the Drs2p P-type ATPase implicated in phospholipid translocation in Saccharomyces cerevisiae. Mol Biol Cell. 2004;15(7):3418-32. doi: 10.1091/mbc.e03-11-0829.

3. Mioka T, Fujimura-Kamada K, Mizugaki N, Kishimoto T, Sano T, Nunome H, et al. Phospholipid flippases and Sfk1p, a novel regulator of phospholipid asymmetry, contribute to low permeability of the plasma membrane. Mol Biol Cell. 2018;29(10):1203-18. doi: 10.1091/mbc.E17-04-0217.

4. Takeda M, Yamagami K, Tanaka K. Role of phosphatidylserine in phospholipid flippase-mediated vesicle transport in Saccharomyces cerevisiae. Eukaryot Cell. 2014;13(3):363-75. doi: 10.1128/EC.00279-13.

5. Zendeh-boodi Z, Yamamoto T, Sakane H, Tanaka K. Identification of a second amphipathic lipid-packing sensor-like motif that contributes to Gcs1p function in the early endosome-to-TGN pathway. J Biochem. 2013;153(6):573-87. doi: 10.1093/jb/mvt025.

6. Miyasaka M, Mioka T, Kishimoto T, Itoh E, Tanaka K. A complex genetic interaction implicates that phospholipid asymmetry and phosphate homeostasis regulate Golgi functions. PLoS One. 2020;15(7):e0236520. doi: 10.1371/journal.pone.0236520.

7. Kishimoto T, Yamamoto T, Tanaka K. Defects in structural integrity of ergosterol

and the Cdc50p-Drs2p putative phospholipid translocase cause accumulation of endocytic membranes, onto which actin patches are assembled in yeast. Mol Biol Cell. 2005;16(12):5592-609. doi: 10.1091/mbc.e05-05-0452.

8. Longtine MS, McKenzie A, 3rd, Demarini DJ, Shah NG, Wach A, Brachat A, et al. Additional modules for versatile and economical PCR-based gene deletion and modification in Saccharomyces cerevisiae. Yeast. 1998;14(10):953-61. doi: 10.1002/(SICI)1097-0061(199807)14:10<953::AID-YEA293>3.0.CO;2-U.

9. Brachmann CB, Davies A, Cost GJ, Caputo E, Li J, Hieter P, et al. Designer deletion strains derived from Saccharomyces cerevisiae S288C: a useful set of strains and plasmids for PCR-mediated gene disruption and other applications. Yeast. 1998;14(2):115-32. doi: 10.1002/(SICI)1097-0061(19980130)14:2<115::AID-YEA204>3.0.CO;2-2.
