## Supplementary material for "Phospholipid flippases and Sfk1 are essential for the retention of ergosterol in the plasma membrane": S2 Table. Plasmids used in this study.

| Plasmid | Characteristics | Reference or source |
|---|---|---|
| YEplac195 | *URA3 2µm* | [1] |
| YEplac181 | *LEU2 2µm* | [1] |
| pRS306 | *URA3* | [2] |
| pRS316 | *URA3 CEN* | [2] |
| pRS315 | *LEU2 CEN* | [2] |
| pBluescript II SK+ | | [3] |
| pKT2210 [pRS316-*SFK1*] | *SFK1 URA3 CEN* | This study |
| pKT2180 [pRS315-*SFK1*] | *SFK1 LEU2 CEN* | This study |
| pKT1252 [pRS315-*LEM3*] | *LEM3 LEU2 CEN* | [4] |
| pKT2211 [pRS315-*CRF1*] | *CRF1 LEU2 CEN* | This study |
| pKT2212 [YEplac195-*OSH2*] | *OSH2 URA3 2µm* | This study |
| pKT2213 [YEplac195-*OSH3*] | *OSH3 URA3 2µm* | This study |
| pKT2175 [YEplac195-*KES1*] | *KES1 URA3 2µm* | This study |
| pKT2114 [YEplac195- $KES1^{E117A}$] | $KES1^{E117A}$ *URA3 2µm* | This study |
| pKT2115 [YEplac195- $KES1^{L111D}$] | $KES1^{L111D}$ *URA3 2µm* | This study |
| pKT2116 [YEplac195- $KES1^{Y97F}$] | $KES1^{Y97F}$ *URA3 2µm* | This study |
| pKT2217 [YEplac195-*OSH5*] | *OSH5 URA3 2µm* | This study |
| pKT2218 [YEplac195-*OSH6*] | *OSH6 URA3 2µm* | This study |
| pKT2219 [YEplac195-*OSH7*] | *OSH7 URA3 2µm* | This study |
| pKT2220 [pRS316-GFP-D4H] | $P_{TPI1}$-*GFP-D4H URA3 CEN* | This study |
| pKT2221 [pRS316-GFPenvy-D4H] | $P_{TPI1}$-*GFPenvy-D4H URA3 CEN* | This study |
| pKT2017 [YEplac181-*KES1*] | *KES1 LEU2 2µm* | This study |
| pKT2222 [pBluescript II SK+-*DNF3-3xGFP*] | *DNF3-Cterminal-3xGFP CaURA3* | This study |
| pKT2223 [pBluescript II SK+-*SFK1-3xGFP*] | *SFK1-Cterminal-3xGFP CaURA3* | This study |
| pKT2224 [pBluescript II SK+-*VPH1-3xGFP*] | *VPH1-Cterminal-3xGFP CaURA3* | This study |
| pKT2225 [pBluescript II SK+-*SFK1-3xmCherry*] | *SFK1-Cterminal-3xmCherry CaURA3* | This study |
| pKT2226 [pRS306-*GFP-evt2-2PH*] | $P_{TPI1}$-*GFP-evt2-2PH URA3* | This study |
| pKT2227 [YEplac181-*SFK1-mCherry*] | *SFK1-mCherry LEU2 2µm* | This study |
| pKT2228 [pRS306-*upc2-1*] | *upc2-1 URA3* | This study |
| pKT2229 [pRS306-*GFP-ER*] | *GFPenvy*-$SCS2^{220\text{-}244}$ *URA3* | This study |
| pKT2230 [YEplac195-*KanMX6*] | *KanMX6 URA3 2µm* | This study |
| pKT2231 [YEplac195-*KES1-KanMX6*] | *KES1 KanMX6 URA3 2µm* | This study |
| pKT2232 [pColdI-GFP-D4H] | GFP-D4H | This study |
| pKT2233 [pColdI-GFPenvy-D4H] | GFPenvy-D4H | This study |

**Reference**

1. Gietz RD, Sugino A. New yeast-Escherichia coli shuttle vectors constructed with in vitro mutagenized yeast genes lacking six-base pair restriction sites. Gene.

1988;74(2):527-34. doi: 10.1016/0378-1119(88)90185-0.

2. Sikorski RS, Hieter P. A system of shuttle vectors and yeast host strains designed for efficient manipulation of DNA in Saccharomyces cerevisiae. Genetics. 1989;122(1):19-27.

3. Alting-Mees MA, Short JM. pBluescript II: gene mapping vectors. Nucleic Acids Res. 1989;17(22):9494. doi: 10.1093/nar/17.22.9494.

4. Kishimoto T, Yamamoto T, Tanaka K. Defects in structural integrity of ergosterol and the Cdc50p-Drs2p putative phospholipid translocase cause accumulation of endocytic membranes, onto which actin patches are assembled in yeast. Mol Biol Cell. 2005;16(12):5592-609. doi: 10.1091/mbc.e05-05-0452.
