## Supplementary figures and images for "Phospholipid flippases and Sfk1 are essential for the retention of ergosterol in the plasma membrane"

### S1 Raw Images. From Figs. 5C, 5D and 7A and S10 Fig.

Fig 5C

GFP-D4H

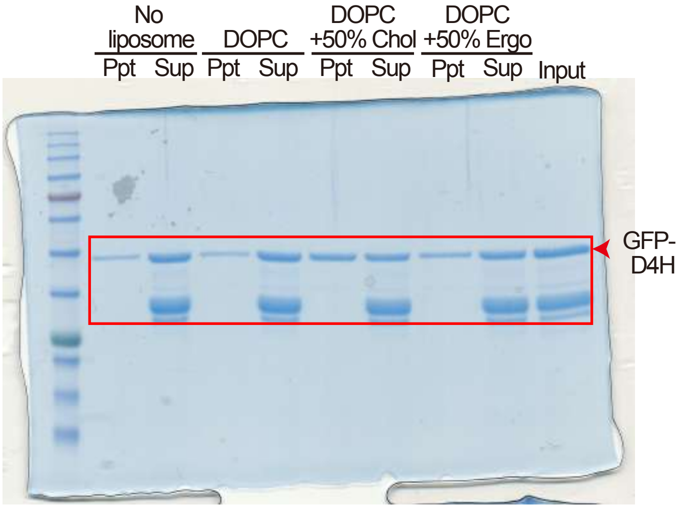


GFPenvy-D4H

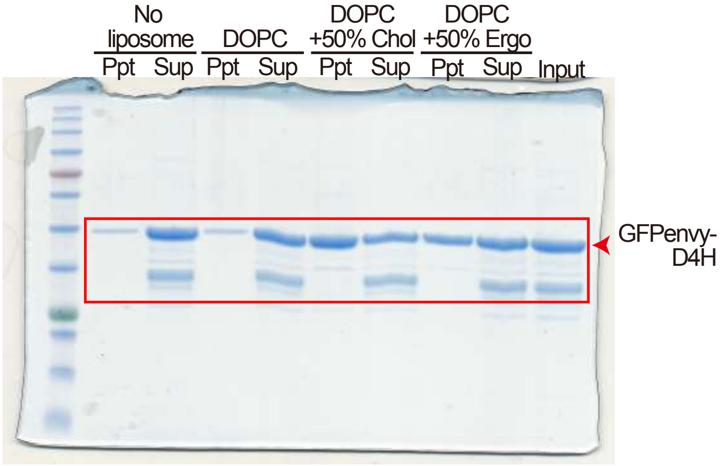


Fig 5D

GFPenvy-D4H

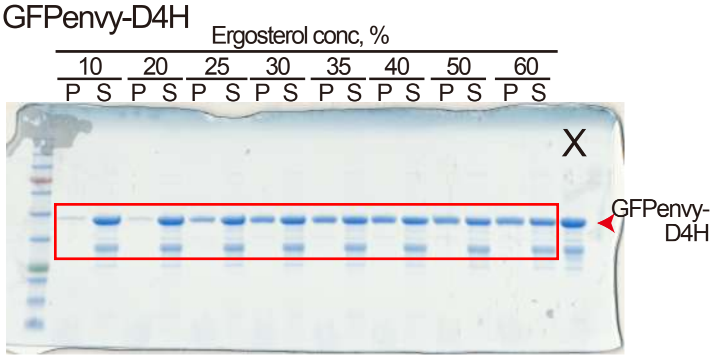

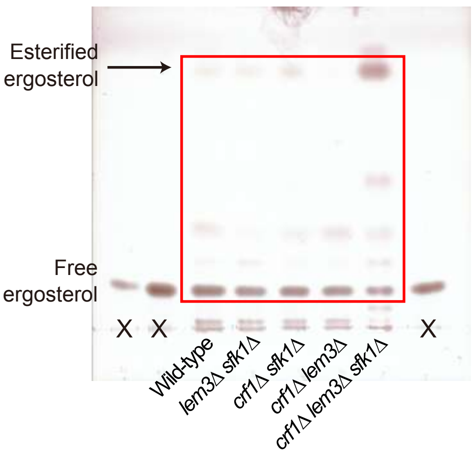

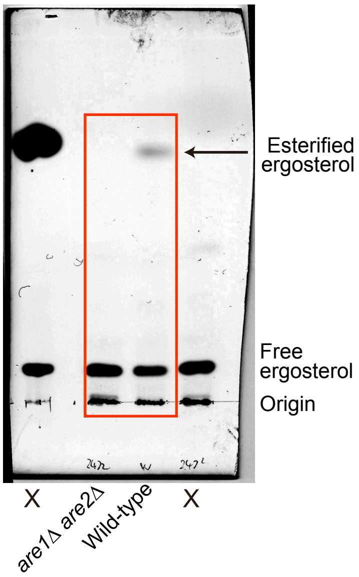
